# Non-enzymatic oxylipin production in a mudflat microphytobenthic biofilm: evidence of a diatom response to light

**DOI:** 10.1101/2024.05.29.596382

**Authors:** Caroline Doose, Camille Oger, Lindsay Mas-Normand, Thierry Durand, Cédric Hubas

**Affiliations:** Muséum National d’Histoire Naturelle, UMR BOREA, MNHN-CNRS-UCN-UPMC-IRD-UA, Station Marine de Concarneau, Concarneau, France; Institut des Biomolécules Max Mousseron, IBMM, Pôle Chimie Balard Recherche, UMR 5247, Université de Montpellier, CNRS, ENSCM, 919 route de Mende, 34293 Montpellier, France

**Keywords:** Microphytobenthos, oxylipins, isprostanoids, light acclimations, Mudflat biofilm

## Abstract

The microphytobenthos (MPB) is a diatom dominated microbial community of primary producers inhabiting the mudflat sediments. In one hand, the benthic diatoms display photo-protective strategies to face extreme light variations susceptible to generate cellular oxidative stress. In the other hand, oxidative stress induces the production of reactive oxygen species (ROS) that generate oxylipins, oxygenated metabolites of polyunsaturated fatty acids (PUFAs), which are among the known chemical mediators in diatoms. However, non-enzymatically generated oxylipins known as isoprostanoids or isofuranoids are poorly studied in diatoms. To better understand the roles of the latter in migrational MPB light response, we investigated the effect of different light irradiances corresponding to dark (D), low light (LL, 50 and 100 µmol. photons. m^−2^. s^−1^PAR), medium light (ML, 250 µmol. photons. m^−2^. s^−1^ PAR), high light (HL, 500, 750 and 1000 µmol. photons. m^−2^. s^−1^PAR), on the isoprostanoids production by the biofilm’s organisms. The PUFAs precursors of the varying oxylipins evidenced a diatoms response to light irradiance. Under 1000 PAR condition, the total amount of isoprotanoids increased, indicating an oxidative stress response. Isoprostanes (IsoPs) and prostaglandins (PGs) characterized the HL conditions and evidenced lipid peroxidation probably linked to the higher generation of ROS by the photosynthesis. On the contrary, the phytoprostanes (PhytoPs) characterized the LL and ML where the de-epoxidation state was low and ROS scavengers probably not overwhelmed. This first investigation of non-enzymatic oxylipin production by a microphytobenthic biofilm under different light irradiances highlighted the interest to explore their potential signaling roles related to MPB light responses.

## 1 Introduction

The microphytobenthos (MPB) is a microbial community of primary producers usually dominated by motile pennate diatoms (Haubois et al., 2005; Méléder et al., 2007; Ribeiro et al., 2013). The MPB sorely contributes to the total primary production of the oceans, representing thus a substantial food source for invertebrates, fish, and wading birds (Beninger and Paterson, 2018; Macintyre et al., 1996; Underwood and Kromkamp, 1999; Werner et al., 2006), and taking an essential role in local socio-economic activities (Lebreton et al., 2019) as well as in the global carbon cycle (Hope et al., 2020) .

The MPB present in mudflats sediments is subjected to strong variability characteristic of the intertidal environments with notably high changes of solar irradiance (Prelle and Karsten, 2022; Woelfel et al., 2014). Diatoms, such as the other photosynthetic organisms, generate reactive oxygen species (ROS) as by-products during the photosynthesis. Under high irradiance exposures, the ROS production of PS II reaction centers can overwhelm the antioxidant systems of the microalgae (Foyer, 2018). Because ROS are highly reactive, they can damage important cells components such as membranes and DNA (Dall’Osto et al., 2010; Havaux and Niyogi, 1999; Krieger-Liszkay et al., 2008; Triantaphylidès and Havaux, 2009). To face to the extreme intertidal light variations, susceptible to generate cellular oxidative stress, benthic diatoms present in muddy sediment habitats display physiological as well as behavioral photo-protective strategies (Barnett et al., 2020; Cartaxana et al., 2011). The physiological responses consist of the dissipation of the energy excess through non-photochemical quenching (NPQ) of chlorophyll (Chl) a fluorescence, the adjustment of light-harvesting pigment production and the ROS detoxification (Lavaud and Goss, 2014; Lepetit et al., 2013; Nymark et al., 2009). The diatom vertical migration is suspected to be a behavioral photoprotective mechanism since they are observed to move downward the sediment under high light irradiances (Consalvey et al., 2004; Jesus et al., 2023). Indeed, it appears to be a diatoms strategy to adapt their vertical positioning to their optimal photon irradiance threshold which also depend on wavelength, and spectral quality of light (Cartaxana et al., 2011; Jesus et al., 2006; Prins et al., 2020; Serôdio et al., 2012).

Given that some studies have indicated the involvement of reactive oxygen species (ROS) in signaling processes in microorganisms (D’Autréaux and Toledano, 2007), and considering their purposeful generation by plants to regulate various metabolic activities such as defense against pathogens, programmed cell death, and stomatal behavior (Apel and Hirt, 2004), our understanding of ROS has evolved in recent decades (Foyer et al., 2017; Noctor and Foyer, 2016). It now appears clear that their roles are diverse and not solely detrimental to cellular functioning.

Because ROS are directly generated during the photosynthesis, are dependent on the efficiency of the latter, as well as the antioxidant systems, they probably play an important role in the induction of photo-protective responses such as NPQ and vertical migration. Despite the intracellular space and chloroplast are highly concentrated in antioxidant, ROS can act locally but also as signaling molecule by being transported to different organelles with maximal distances ranging from 1 nm for ^•^OH to more than 1 µm for the hydrogen peroxide (H_2_O_2_) (Dumanović et al., 2021; Knieper et al., 2023; Mittler, 2017). The oxidation by-products of ROS detoxification molecules such as oxidized glutathione (Meyer, 2008) and those given by ^1^O_2_ and carotenoids also can give rise to signaling molecules (Ramel et al., 2012). These by-products can modulate various biological processes, including transcription, post-translational modification, and protein–protein interactions, notably by impacting the oxidation state of thiol groups in redox-sensitive proteins (Dietz, 2008; Meyer, 2008).

The oxylipins are also byproducts generated from ROS reaction from polyunsaturated fatty acids (PUFAs), called lipid peroxidation (Améras et al., 2003; Jahn et al., 2008; Triantaphylidès et al., 2008). These compounds represent the best described signaling molecules in diatoms (Orefice et al., 2022; Ruocco et al., 2020). The oxylipins can influence other species abundance and fitness through the antipredator, antibacterial, infochemical and allelochemical functions of oxylipins (Meyer et al., 2018; Ruocco et al., 2020; Russo et al., 2020). They can be produced by several enzymatic and non-enzymatic processes, giving an important structures’ diversity (Galano et al., 2017; Gerwick et al., 1991; Longini et al., 2017). The enzymatic lipoxygenase pathways have been shown to be species dependent in the marine diatom’s genus *Pseudo-nitzschia (Lamari et al., 2013)*, while the oxylipin structure from non-enzymatic process, named isoprostanoids, depend only on the ROS reaction within a bis-allylic position of PUFAs’ double-bonds in the cells (Galano et al., 2017). The non-enzymatic oxylipins are likely less specific to species, making isoprostanoids promising candidates for transmitting signals between various kingdoms of organisms present in the microphytobenthic biofilm. In addition, previous works showed that the presence of H_2_O_2_ and copper in culture media induced C18-, C20- and C22-derived isoprostanoid production changes in several diatoms and other microalgae species and some of them were observed to trigger biological responses (Linares-Maurizi et al., 2023; Lupette et al., 2018; Vigor et al., 2020).These oxylipins could be thus involved in the MPB responses to physiological changes and environmental variations such as light exposure.

The oxylipins biosynthesis by microalgae, especially diatoms, started to trigger a lot of interest in the last few years (Di Dato et al., 2020a, 2020b, 2019) but little emphasis has been done on non-enzymatic pathways (Orefice et al., 2022; Vigor et al., 2020). Also, their production in microphytobenthic biofilm was never studied. Thus, to better understand the roles of the latter in migrational MPB light response, we investigated the effect of different light irradiances corresponding to dark (D), low light (LL, 50 and 100 µmol photons m^−2^ s^−1^PAR), medium light (ML, 250 µmol photons m^−2^ s^−1^PAR), high light (HL, 500, 750 and 1000 µmol photons m^−2^ s^−1^PAR) on their presence in the biofilm’s organisms.

## 2 Materials and methods

### 2.1 Biofilm sampling and light exposure

The Biofilm’s sample used to carry this study were the same as those generated after 30 min of light exposure in (Doose and Hubas, 2024). The first 2 cm of sediment present in an empty breeding pond of the Marine Station of Concarneau (France; 47◦52.5804’N; 3◦55.026’W) at low tide was sampled to collect the MPB biofilm.

After 24h in half obscurity (PAR < 6 µmol. photons. m^−2^. s^−1^PAR) to let it settle, the top 5 mm of the sediment containing the MPB was sampled and re-suspended in 250 ml of filtered seawater. A volume of 6 mL of resuspended biofilm was added in 5 cm diameter Petri dishes and left 24h in half obscurity (PAR < 6 µmol. photons. m^−2^. s^−1^PAR) to let it settle. A number of 5 Petri dishes (n=5) were then placed under dark (D), 50, 100 (LL), 250 (ML), 500, 750 and 1000 (HL) µmol. photons. m^−2^. s^−1^PAR where light was generated by Led Lights (SL 3500, white warm, Photon Systems Instruments). After 30 min of exposure, liquid nitrogen was poured in the Petri dishes to freeze immediately the sediment without disturbance. The samples were freeze-dried and stored at -80°C waiting for the subsequent analysis.

### 2.2 Oxylipins

#### 2.2.1 Sample preparation

Non-enzymatic oxylipins were extracted using a protocol that was previously published in prior work on marine macroalgae (Vigor et al., 2018) with some modifications. For the extraction, 150 mg of fresh biomass were placed in lysing matrix tubes (lysing matrix D, MP Biochemicals, Illkirch, France) with 25 µL of BHT (butylated hydroxytoluene 1% in methanol), 1 mL of H_2_O (HPLC grade) and 4 µL of Internal Standards Mixture (ISM n°18) (1 ng/µL). The sample was then grinded using a FastPrep-24 (MP Biochemicals) at a speed of 6.5 m/s for 30 s. The mixture was transferred into a 15 mL falcon tube with 3 x 1 mL of cold chloroform/methanol mixture (2:1) and was stirred with a vortex mixer for 30 s between each transfer. A volume of 0.5 mL of phosphate buffer (50 mM, pH 2, prepared with NaH_2_PO_4_ and H_3_PO_4_) saturated in NaCl and stirred with a vortex mixer for 30 s) was added to the mixture. Then, 3 mL of cold chloroform/methanol mixture (2:1) was added and stirred with a vortex for 30 s. The samples were then centrifuged at 4000 rpm for 5 min at 4°C. The lower organic phase was collected in Pyrex tubes and was then dried using SpeedVac apparatus, at 60°C for 1 hour.

To extract lipid fraction, the dried extract was hydrolyzed by adding 950 µL of 1 M KOH and was incubated at 40°C for 30 min with a vertical rotator (100 rpm). The mixture was added with 1 mL of 40 mM formic acid prior to starting the solid phase extraction. After that, samples were loaded on preconditioned Oasis mixed polymeric sorbent cartridges (Oasis MAX Cartridge, 60 mg, Waters). The undesired compound was then eliminated using 2 mL of NH_3_ 2% (v/v), 2mL of MeOH/20 mM formic acid (30:70; v/v), 2 mL of hexane, and 2 mL of hexane/ethyl acetate (70:30; v/v). Finally, isoprostanoids/isofuranoids/PG were eluted by adding 2 x 1 mL of a mixture of Hexane/EtOH/ Acetic acid (70:29.4:0.6; v/v/v). The samples were dried using SpeedVac at 60°C for an average 1 h.

The dried extracts were reconstituted with 100 µL of mobile phase solvents (H_2_O/ACN; 83:17; v/v) and then stirred with a vortex, ultrasounds 2 minutes and vortex and later filtered in 0.45 µm Eppendorf (Nanosep Centrifugal Devices) with a centrifugation at 10 000 rpm for 1 min at room temperature. A volume of 80 µL was transferred in an HPLC analytic vial for further analysis, and the remaining 20 µL were transferred in another HLPC vial for spiking QC. Note that for the QC, a volume of 4 µL of Prostamix GR57 SM0.5 containing all oxylipin standards at 0.5 ng/µL. The analysis was completed by injecting 5 µL of the extract into the micro-LC-MS/MS 5500 QTrap system, which uses high-performance liquid chromatography coupled to tandem mass spectrometry.

#### 2.2.2 Quantification measurements by micro-LC-MS/MS

An Eksigent micro-High performance liquid chromatography (HPLC) 200 Plus (Sciex Applied Biosystems, Framingham, MA, USA) equipped with CTC Analytics AG (Zwingen, Switzerland) was used and all analyses were carried out on a HALO C18 analytical column (100 x 0.5 mm, 2.7 µm; Eksigent Technologies, CA, USA) maintained at 40°C. The mobile phases consisted of a binary gradient of H_2_O with 0.1% (v/v) HCO_2_H (solvent A) and ACN/ MeOH 80:20 (v/v) (solvent B) with a flow rate of 0.03 mL.min^-^1 and an injection volume of 5 µL. The elution gradient was as follows: 17% B at 0 min, 17% B at 2.6 min, 21% B at 2.85 min, 25% B at 7.3 min, 28.5% B at 8.8 min; 33.3% B at 11 min; 40% B at 15 min, 95% B at 16.5 min for 1.5 min.

Using an electrospray ionization (ESI) in negative mode, mass spectrometry analyses were performed on an AB Sciex QTRAP 5500 (Sciex Applied Biosystems, ON, Canada). The source is maintained at −4.5 kV, and nitrogen flow serves as curtain gas at 30 psi and a nebulization assist at 20 psi, at room temperature.

In order to analyze the targeted compounds with a detection window of 90 s, the monitoring of the ionic fragmentation products of each deprotonated analyte [M-H]^-^ molecule was carried out in Multiple Ion Monitoring (MRM) detection mode using nitrogen as the collision gas. Two transitions for quantification (T1) and specification (T2) were predetermined by MS/MS analysis of standards. LC-MS/MS data acquisition was performed using the Analyst® software (Sciex Applied Biosystems), which drives the mass spectrometer. The peak integration and quantification of analytes were processed by MultiQuant 3.0 software (Sciex Applied Biosystems).

Out of the 54 oxylipins’ standards in hands from several omega 3 and omega 6 PUFAs, eight PhytoPs and two PhytoFs from α-linolenic acid (ALA), six IsoPs and one PG from eicosapentaenoic acid (EPA), four IsoPs and one PG from arachidonic acid (AA), one dihomo-IsoP from adrenic acid (AdA), and seven neuroprostanes from docosahexaenoic acid (DHA) for the non-enzymatic oxylipins, and three prostaglandins were highlighted here (see results part).

### 2.3 Data treatment and statistical analysis

The mean of de-epoxydation state data were calculated per light group (LL, 50-100 PAR; ML, 250 PAR; and HL, 500,750 and 1000 PAR). The object generated by a Multiple Factor Analysis MFA performed on oxylipin measurement data was used to run a Between Class analysis on R under the package ade4 following the original script available on Github (https://github.com/Hubas-prog/BC-MFA). The result is named BC-MFA. The normality of the pigment and oxylipin data and their residual distribution was tested using the Shapiro–Wilk test. When residuals followed a normal distribution, a one-way ANOVA was performed to detect significant light effects on oxylipin concentrations in the biofilm. When normality was not verified, the non-parametric Van der Waerden test was performed with the R package agricolae. Outliers were tested using the 1.5 times the interquartile range (IQR) rule. The test of correlation was performed on R with the Pearson method.

## 3 Results

### 3.1 The effect of light on oxylipin fingerprint in biofilm

The LC-MS targeted analysis allowed to measure and quantify 29 oxylipins (supplementary materials, S1) from 5 different families: neuroprostanes (NeuroPs), isoprostranes (IsoPs), phytoprostanes (PhytoPs), phytofuranes (PhytoFs) and prostaglandins (PG). The BC-MFA analysis presented in the Fig.1 shows the partitioning of MPB samples depending on their amount of oxylipin measured. The total inertia of the dataset explained (T.I.E) by the light treatments was 37 %. The first dimension of the BC-MFA scoreplot distinguished the dark from the 1000 PAR treatment as well as the LL and ML from the HL treatments. The dark, LL and ML treatments were characterized by oxylipins from the PhytoP family while the HL were characterized by oxylipins from the IsoP and PG families. This analysis allowed to identify 12 compounds (out of 29) significantly influenced by this light gradient. The production of molecules forms the NeuroP and the PhytoF families were not affected by the light irradiances.

**Fig 1.**
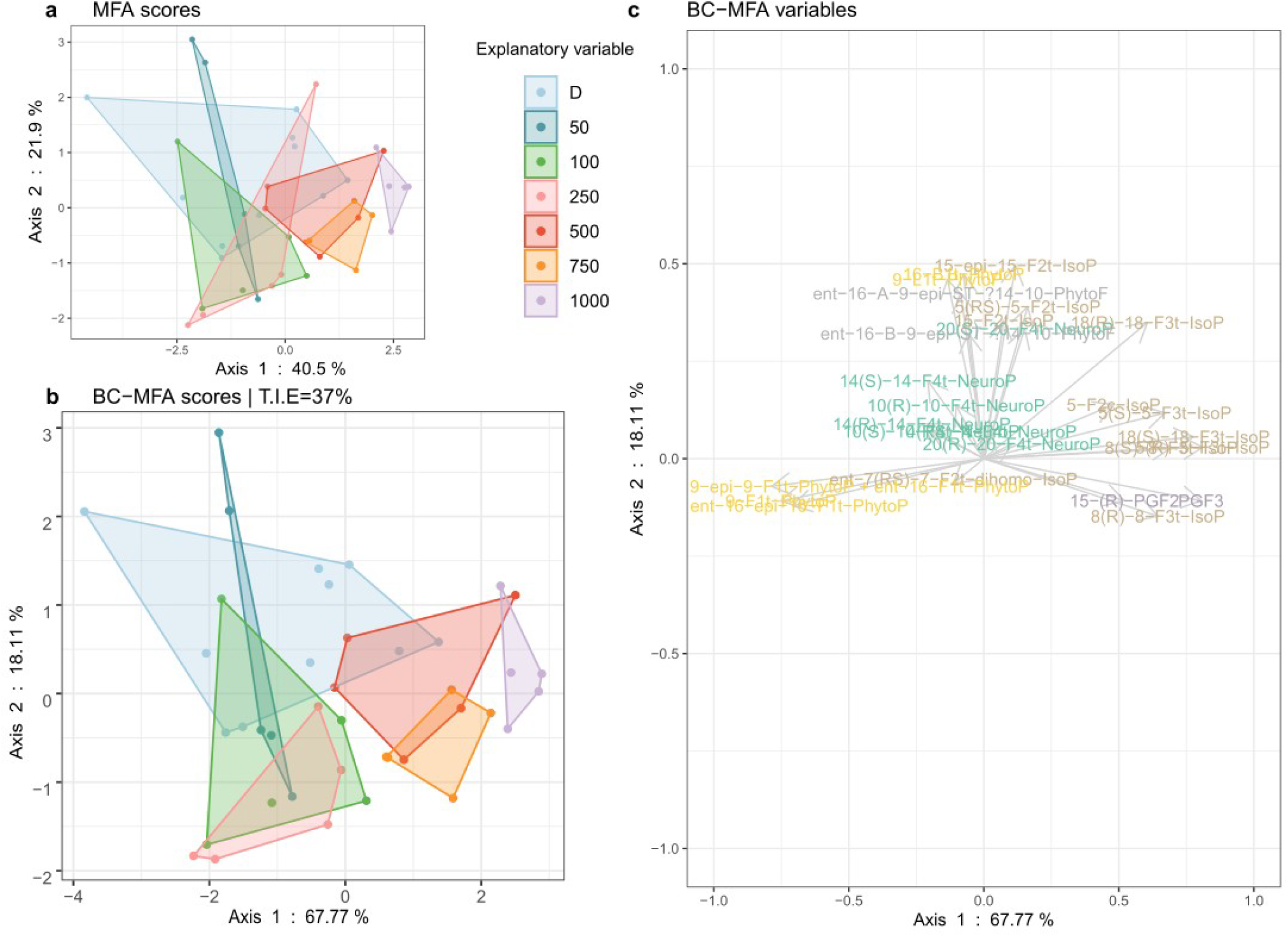
Between Class (BC) analysis realized on a Multiple Factor Analysis (MFA) object (BC-MFA) on oxylipins measured in biofilm exposed to dark (D, n=10) and to irradiances of low lights (LL: 50, 100), medium light (ML: 250) and high lights (HL: 500, 750, 1000) in µmol. photons. m^−2^. s^−1^ PAR (n=5).

The highest total amount of oxylipins presented in Fig. 2 was measured in biofilm exposed to the 1000 PAR treatment with 484 ± 27 pg/mg dw. The lowest values were observed in biofilm under dark (383 ± 39 pg/mg dw), 250 and 500 PAR (385 ± 17 pg/mg dw for both). Significant differences were observed between biofilm under the 1000 PAR treatment and all the other irradiances, except for the 100 PAR conditions (ANOVA, p< 0.05). The major oxylipins, constituting more than 5% of the total oxylipin measured in the biofilm, included 18-*epi*-18-F_3t_-IsoP (EPA), 5-*epi*-5-F_3t_-IsoP (EPA), 5-F_3t_-IsoP, 9-*epi*-9-F_1t_-PhytoP + *ent*-16-F_1t_-PhytoP, 9-F_1t_-PhytoP, and PGF_3_.

**Fig 2.**
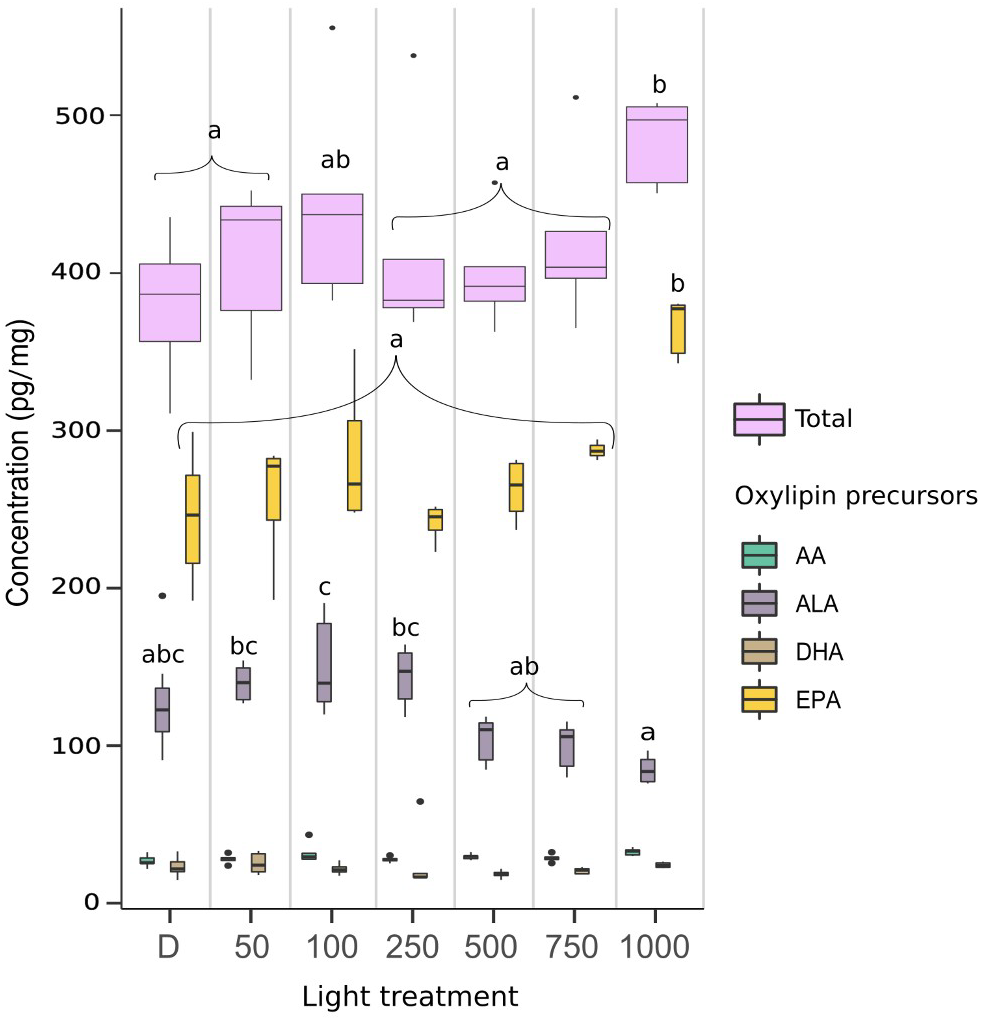
Amount of oxylipins in pg/mg dw per fatty acid precursors and in total measured in biofilm exposed to dark (D, n=10), and to irradiances of low lights (LL: 50, 100), medium light (ML: 250) and high lights (HL: 500, 750, 1000) in µmol photons m^−2^ s^−1^PAR (n=5). AA (arachidonic acid), AdA (adrenic acid), ALA (α-linolenic acid), DHA (docosahexaenoic acid), EPA (eicosapentaenoic acid). Outliers were identified only in the total oxylipins data using the IQR for the 100, 250, 500 and 750 PAR conditions (for which n=4). A one-way ANOVA was performed on the data to detect significant differences between treatments, as indicated by letters (p < 0.05).

### 3.2 Oxylipins variation in function of their PUFAs precusors

The Fig 3. show the proportion of the metabolite’s precursor. The oxylipins measured in the biofilm were mainly derive from the eicosapentaenoic acid (EPA) (Guy et al., 2014; Morrow et al., 1990) representing more than 60% of the total amount. The total of α-linolenic acid (ALA) metabolites (Parchmann and Mueller, 1998) represented more than 10%, the arachidonic acid (AA) (Guy et al., 2014; Morrow et al., 1990) and the docosahexaenoic acid (DHA) (Nourooz-Zadeh et al., 1998; Roberts et al., 1998) represented around 7 and 5 % respectively. One metabolite measured was derived from the adrenic acid (AdA), this latter represents therefore less than 1% of the total FA precursor. Only the total of EPA and ALA metabolites varied under the different light irradiation. The total EPA metabolites increased with increasing irradiances from 250 PAR to the 1000 PAR conditions. Conversely, the total ALA metabolites followed the exact opposite pattern, with the highest value under the 250 PAR condition. The 750 and 1000 PAR conditions are significantly lower than the dark, LL, and ML conditions. The total amount of oxylipins per FA precursor, presented in Fig. 2, showed significantly higher amounts of oxylipins derived from EPA under the 1000 PAR irradiance than in the other light conditions. Conversely, the amount of oxylipins derived from ALA was significantly higher under the 100 PAR irradiances than under HL. The variations in EPA and ALA derived oxylipin percentage are thus explained by both the increase of EPA-derived compounds and the decrease of ALA-derived compounds.

**Fig 3.**
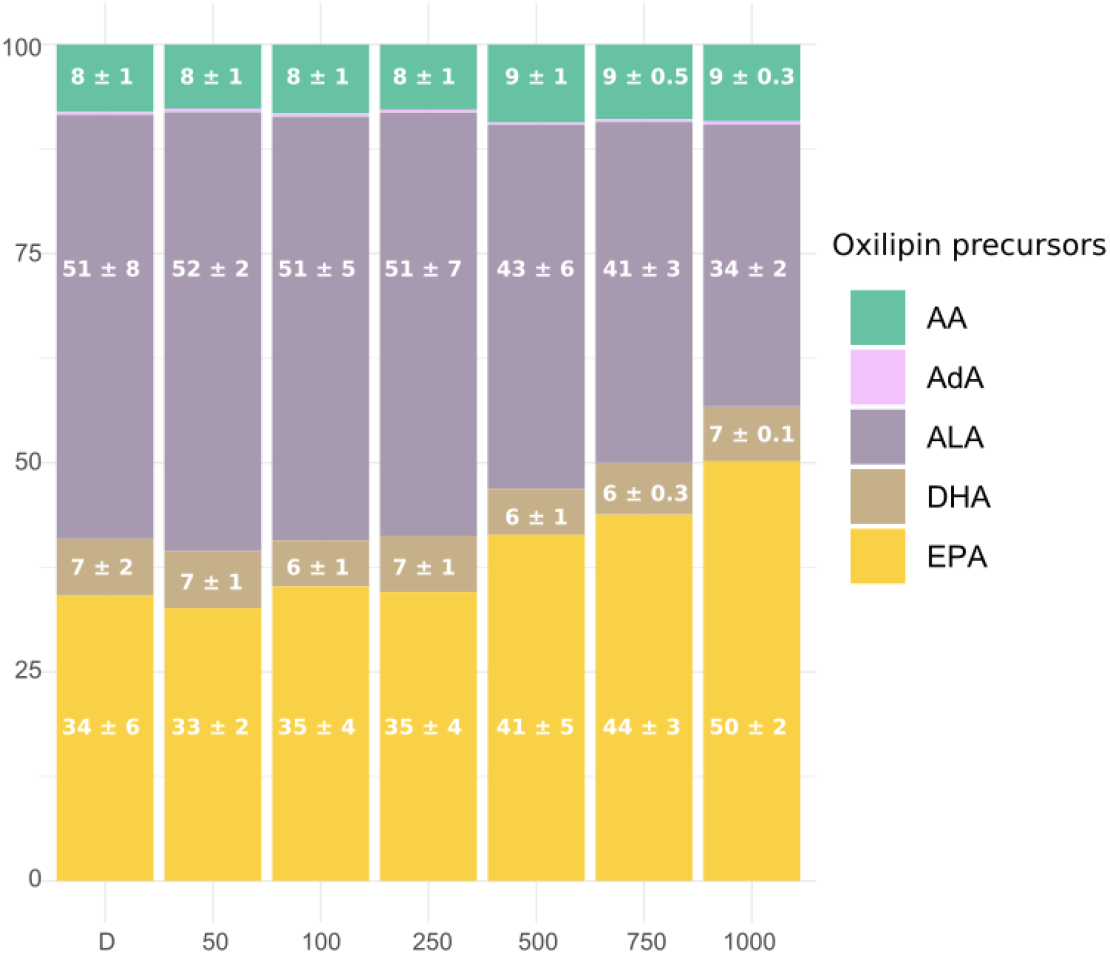
Relative abundance distribution in percentage of oxidated polyunsaturated fatty acids (NEO-PUFAs) in the MPB biofilm: AA (arachidonic acid), AdA (adrenic acid, <1%), ALA (α-linolenic acid), DHA (docosahexaenoic acid), EPA (eicosapentaenoic acid). Significant differences between light exposure are indicated by letters (ANOVA, p < 0.05; n = 5)

The Fig 4. presents the values of oxylipins which significantly varied between the different light conditions. Half of them are derived from the EPA. The amount of the 6 EPA-derived oxylipins and the 2 AA-derivatives which varied under the light were all significantly higher under 1000 PAR than in the dark condition (ANOVA, p< 0.05). However, significant differences were also observed between the 1000 PAR and LL and ML conditions notably for the 18-*epi*-18-F_3t_-IsoP and the 5-*epi*-5-F_3t_-IsoP the LL.

**Fig 4.**
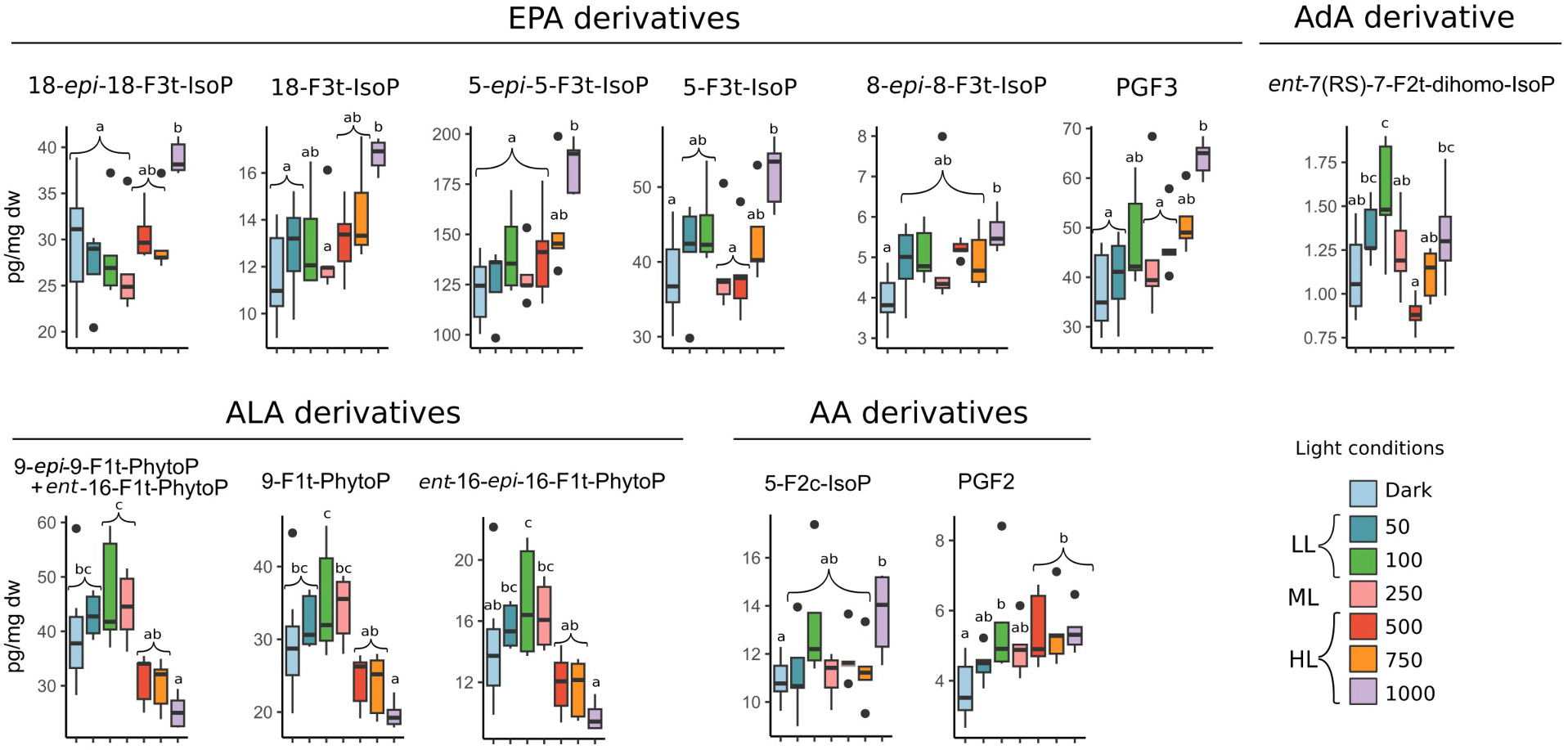
Concentration of the oxylipins in pg/mg dw which significantly varied in biofilm exposed to dark (D, n=10) and to irradiances of low lights (LL: 50, 100), medium light (ML: 250) and high lights (HL: 500, 750, 1000) in µmol photons m^−2^ s^−1^PAR. A one-way ANOVA was performed on the data to detect significant differences between treatments, as indicated by letters (p < 0.05; n = 5). The oxylipins were gathered in groups corresponding to their fatty acid precursors AA (arachidonic acid), AdA (adrenic acid), ALA (α-linolenic acid), DHA (docosahexaenoic acid), EPA (eicosapentaenoic acid).

The oxylipins derived from ALA exhibited variations exclusively within the F_1t_-series, and followed an inverse trend to that observed in EPA metabolites. The amount of ALA-derived oxylipins reached a maximum under 100 PAR with 47 ± 10 pg/mg dw for the 9-*epi*-9-F_1t_-PhytoP + *ent*-16-F_1t_-PhytoP and decrease following the increasing irradiance level to reach a minimum value under 1000 PAR with 8 ± 1 pg/mg dw for the *ent*-16-*epi*-16-F_1t_-PhytoP.

The only AdA metabolite measured in this analysis was the *ent*-7(*RS*)-7-F_2t_-dihomo-IsoP. Its amount increased following the increasing light irradiances between the dark and the 100 PAR conditions as well as between the 500 PAR and the 1000 PAR condition and decreasing between the 100 PAR and the 500 PAR conditions. The maximum amount was observed under 100 PAR with 1.5 ± 0.3 pg/mg dw which was significantly higher than the values observed under the dark, 250, 500 and 750 PAR conditions (ANOVA, p< 0.05). The amount measured under the 1000 PAR condition were also significantly higher than those found the 500 PAR condition (ANOVA, p< 0.05).

All the oxylipin amount significantly varying under the different light condition were correlated with the irradiance’s level except for the *ent*-7(*RS*)-7-F_2t_-dihomo-IsoP (table 1). The PhytoPs derived from ALA were negatively correlated with the increase of photon flux. In the contrary, the oxylipin amount derived from EPA and AA were positively correlated with the light increase. The strongest correlation was found for the 5-*epi*-5-F_3t_-IsoP which is also the most abundant compound measured.

**Table 1.** Significant correlation between the light irradiances levels and the amount in oxylipins measured in the biofilm.

| Oxylipins | Fatty acid precursor | P value | Correlation coefficient |
| --- | --- | --- | --- |
| 15-(R)-PGF<sub>2</sub> | AA | 8,8E-04 | 0,51 |
| 5-F<sub>2c</sub>-IsoP |  | 4,5E-02 | 0,32 |
| PGF<sub>2</sub> |  | 5,7E-03 | 0,43 |
| 18-*epi*-18-F<sub>3t</sub>-IsoP | EPA | 1,7E-03 | 0,48 |
| 18-F<sub>3t</sub>-IsoP |  | 1,6E-05 | 0,63 |
| 5-*epi*-5-F<sub>3t</sub>-IsoP |  | 4,8E-07 | 0,70 |
| 5-F<sub>3t</sub>-IsoP |  | 2,9E-03 | 0,46 |
| 8-*epi*-8-F<sub>3t</sub>-IsoP |  | 5,8E-03 | 0,43 |
| 8-F<sub>3t</sub>-IsoP |  | 7,3E-04 | 0,51 |
| PGF<sub>3</sub> |  | 2,0E-06 | 0,67 |
| 9-*epi*-9-F<sub>1t</sub>-PhytoP + *ent*-16-F<sub>1t</sub>-PhytoP | ALA | 8,0E-06 | -0,64 |
| 9-F<sub>1t</sub>-PhytoP |  | 4,7E-05 | -0,60 |
| *ent*-16-*epi*-16-F<sub>1t</sub>-PhytoP |  | 1,1E-04 | -0,57 |

## 4 Discussion

### 4.1 Oxylipin origins regarding their PUFA precursors

The microphytobenthic biofilm encompasses a wide diversity of microorganism taxa, with diatoms overwhelmingly dominating. The isoprostanoids measured in this work depend only on the ROS reaction with the PUFAs and their release as free oxidized lipids by the phospholipases A2 in the cells (Ibrahim et al., 2011; Mallick and Mohn, 2000; Morrow et al., 1992; Roy et al., 2017). Their variation in amount and composition under the different irradiances evidence that the light levels affect ROS production in the microphythobentic biofilm. In the latter, the primary metabolism susceptible to generate rapid and substantial variations in ROS production under light irradiation changes is photosynthesis, the chloroplast of these microorganisms being a major site of ROS generation (Pitzschke et al., 2006). Since FAs are widely recognized as biomarkers (Kelly and Scheibling, 2012; Parrish, 2013), associations with specific biological compartments within the biofilm can be established through the PUFA precursors of light-varying oxylipins. The IsoPs emerged as the predominant varying oxylipins, with EPA identified as their precursor, a well-known diatom marker (Kelly and Scheibling, 2012; Parrish, 2013). The second group of oxylipins that exhibited significant variation under different light irradiances was PhytoPs. They are produced from ALA, which is known to be present in higher concentrations in green algae than in diatoms (D’Souza and Loneragan, 1999; Kelly and Scheibling, 2012). PhytoPs were however identified as the primary non-enzymatic oxylipins produced by the diatom *P. tricornutum*, being measured in the same concentration range as the non-oxidized ALA (around 400 pmol per 1 million cells) (Lupette et al., 2018; Vigor et al., 2020). Therefore, the presence of PhytoPs in the biofilm could also be linked to diatom response.

### 4.2 Non-enzymatic oxylipins evidence distinct oxidative responses to the different light levels

The PUFA precursors implied in oxylipins variation were not the same depending on the light irradiances. The ALA-derived oxylipins were in higher proportion in the dark, LL and ML conditions than under HL and inversely the AA and EPA-derived oxylipins were measured in higher proportion under HL than the other light conditions. The different light irradiance levels appear to trigger peroxidation on specific PUFAs. Moreover, the ALA peroxidation was observed through the variation of the F_1t_-PhytoPs. But the B_1t_-PhytoP and PhytoFs, also derived from this PUFA, didn’t varied. In a recent study, the F_1t_-PhytoPs were also the most abundant series which varied under H_2_O_2_ oxidative stress *P. tricornutum* while the *ent*-16-B_1t_-PhytoP and 16(*RS*)-16 A_1_-PhytoPs measured in low levels didn’t vary (Lupette et al., 2018). Along the specific PUFA peroxidation, the ROS produced under different light conditions may also exhibit preferences for specific reaction site and pathways.

Interestingly, the increase of oxidation induced by light-triggered ROS production did not appear to affect all PUFA precursors. Specifically, in the case of DHA, the peroxidation products, known as NeuroPs, remained constant across varying light irradiances. In related studies, observations in the diatom *C. gracilis (Vigor et al., 2020)* and *P. tricornutum* indicated that NeuroPs did not vary under H_2_O_2_ oxidative stress. Moreover, these compounds were found to be produced in the same concentration range as their non-oxidized precursors (Lupette et al., 2018). Indeed, ROS are constantly produced as byproducts of metabolisms, such as photosynthesis or photorespiration, which continuously lead to lipid peroxidation by healthy organisms (Knieper et al., 2023; Mueller, 2004).This could partly explain this basal presence of the NeuroPs and the other non-varying oxylipins measured in the microphytobenthic biofilm and under non-stressing light levels.

### 4.3 Light-dependent variations in oxylipin profiles and photosynthesis ROS production

The total of varying EPA-derived oxylipins significantly increase from 750 PAR (Fig.3) to 1000 PAR. Their amount were significantly correlated with the increasing irradiance level. One of them, the 5(*R*)-5-F_3t_-IsoP was the most abundant with values comprised between 100 and 200 pg/mg dw. In the diatom *C. gracilis*, the most abundant isoprostanoids identified were also derived from this PUFA and the 5(*RS*)-5-F_3t_-IsoP accounted for approximately 42% (1.1 µg/g) of the total oxylipins (Vigor et al., 2020). The 8-F_3t_-IsoP or 8-*epi*-8-F_3t_-IsoP was also observed to significantly increase in *P. tricornutum* after 48h under 0.75 mM of H_2_O_2_ (Lupette et al., 2018). But the inverse trend was found after 24h under 1 mM of H_2_O_2_ for the same diatom and C. *gracilis (Vigor et al., 2020)*.

During photosynthesis, H_2_O_2_ is generated notably through the water-water cycle (WWC) (Asada, 1999). In microalgae and cyanobacteria, the WWC is a significant pathway for dissipating excitation energy, accounting for up to 49% of the total electron flux in diatoms (Curien et al., 2016; Waring et al., 2010). The ROS production under high light can exceed the rate of the WWC reactions leading to H_2_O_2_ increase in the chloroplast. It is challenging to determine whether the concentrations of H_2_O_2_ examined in the aforementioned studies are comparable to what photosynthesis might induce under HL conditions in microphytobenthic organisms. This difficulty arises due to uncertainties about the extent of H_2_O_2_ entry into the cell through aquaporins (Knieper et al., 2023; Vogelsang and Dietz, 2022) and the possibility that exposure via the culture medium could induce peroxidation at other membrane sites than those associated with H_2_O_2_ production in the chloroplast under HL. However, significant accumulation of H_2_O_2_ in the diatom *N. epithemioides* was observed in the similar light exposure (30 and 40 min at 1000 µmol photons m^−2^ s^−1^) reaching values between 1 and 1.5 µmol/µg Chl*a* (Waring et al., 2010). This suggest that the observed increase in AA and EPA-derived oxylipins in the biofilm under 1000 PAR might be partly attributed to the heightened production of H_2_O_2_ through photosynthesis in the MPB, particularly in diatoms.

ROS can engage in diverse signaling pathways contingent upon their reactivity and if light stress persists or intensifies, the diversity of generated ROS (H_2_O_2_, ^1^O_2_, O_2_^•−^, etc.) also increases (Foyer, 2018; Waring et al., 2010). The ^1^O_2_, characterized by high reactivity, can initiate signals but not transport them. In contrast, the lower reactivity of H_2_O_2_ allows it to interact with various biological sites, to function as a mobile messenger and to be excreted in the extracellular space (Mullineaux et al., 2018; Schneider et al., 2016). Changes in the amount and composition of ROS under varying levels of light stress at the photosystem sites may partly explain the differences observed in oxylipin production at different irradiance’s levels. Interestingly, the total amount of measured oxylipins did not vary between all the light conditions except for the 1000 PAR condition. This suggest that the peroxidation rate was the same under these first mentioned light irradiances and that amount of ROS could have been maintained at a steady state by the anti-oxidant systems of the cells. The qualitative difference between the LL, ML and the concerned HL (500 and 750 PAR) could be attributed to a different cellular location involved in these oxylipins synthesis as well as different composition of ROS produced under these irradiances. The increase in IsoPs and PGFs observed under 1000 PAR conditions could thus be triggered by specific ROS signature produced under conditions of overwhelmed ROS scavengers.

The increase in PhytoPs was observed under 100 and 250 PAR. Under these light conditions, photosynthesis is presumed to be efficient, considering the Ek and Eopt values measured in the same biofilm samples (Doose and Hubas, 2024), and ROS scavengers are likely not overwhelmed. The increase of these oxylipins triggered under an efficient state of photosynthesis might also suggest that the peroxidation could be attributed to ROS originating from non-photosynthetic organisms. ROS are ubiquitous in marine environments (Diaz et al., 2016; Paul Hansard et al., 2010; Roe et al., 2016; Rose et al., 2008), and bacteria present in natural water are known to contribute to the H_2_O_2_ source (Dixon et al., 2013; Marsico et al., 2015; Vermilyea et al., 2010; Zhang et al., 2016) and probably to O_2_^•−^ source as well (Diaz et al., 2013; Hansel et al., 2019; Learman et al., 2011; Sutherland et al., 2019; Zhang et al., 2016). However, their concentrations in marine environments range from picomolar to hundreds of nanomolar (Diaz et al., 2016; Paul Hansard et al., 2010; Roe et al., 2016; Rose et al., 2008; Rusak et al., 2011). As discussed previously, PhytoPs are very likely produced by diatoms for which only certain glycerolipids (phosphatidylcholine and diacylglyceryl-hydroxymethyl-N,N,N-trimethyl-β-alanine) present in intracellular endomembranes, which contained sufficient amounts of its PUFA precursor (ALA) to justify a role for the production of PhytoPs (Leblond et al., 2013; Lupette et al., 2018). Even if higher concentrations of extracellular ROS are often measured in high-productivity waters and shallow coastal environments (Diaz et al., 2016; Paul Hansard et al., 2010; Roe et al., 2016; Rose et al., 2008; Rusak et al., 2011), their concentrations might be not sufficient to trigger all the intracellular peroxidation in diatoms providing the oxylipins amounts measured in the present study.

The observed increase in PhytoPs may suggest a controlled and regulated response by some microphytobenthic organisms, notably diatoms as discussed above. This could result from peroxidation triggered by a specific ROS signature of photosynthesis in a well-maintained state. Indeed, the cell capacity to detoxify, scavenge, or buffer ROS is suspected to control the quantity and spatial accumulation of ROS, which can be specific to an intracellular site such as a membrane patch or organelle (Knieper et al., 2023; Mittler et al., 2011). Moreover, the antioxidant controls the termination of the peroxidation reaction (Montuschi et al., 2004) and photoprotectants such as xanthophylls are known to have an effect on oxylipins presence (Demmig-adams et al., 2012).This spatial specificity of ROS and antioxidant presence could thus also explain the differences in the distinct oxylipin responses under the different light levels. Moreover, regarding the localisation of the PUFAs precursors in diatom cell, lipid peroxidation might predominantly occur in endomembranes under LL and ML conditions (Lupette et al., 2018), whereas under HL conditions, it might happen in chloroplasts since EPA is known to be a major fatty acid in diatom thylakoïd membranes (Büchel et al., 2022).

### 4.4 EPA and ALA derivatives presence followed the photoacclimation state of MPB

Under 100 PAR, the total amount of oxylipins measured in the biofilm tended to increase and had the highest amount of ALA derivatives, as well as the AdA derivative *ent*-7(*RS*)-7-F_2t_-dihomo-IsoP. However, regarding the oxylipin proportion presented in Fig. 2, the higher and lower percentages for ALA-derived and EPA-derived oxylipins, respectively, were observed under 250 PAR. Interestingly, the Ek value is situated between 100 and 250 PAR, with 187 ± 22 PAR (Doose and Hubas, 2024). This value is suspected to represent the theoretical optimal light level for the MPB and it was suggested that diatoms adjust through their vertical migration in the sediment the irradiance they receive around this Ek value (Jesus et al., 2023). Moreover, a downward migration of the MPB was induced from 250 PAR (visual observation), corresponding to the range of irradiance found in literature to induce mudflat MPB downward movement (Ezequiel et al., 2015; Laviale et al., 2016; Perkins et al., 2010; Serôdio et al., 2008, 2006). The PhytoP synthesis was thus concomitant with the presence of MPB at the sediment surface, thus more exposed to oxidative conditions compare to the anoxic conditions in the sediment. But as discussed previously, the water amounts of ROS are likely not enough to explain the entire presence of these isoprostanoids in the biofilm. Regarding the high values of those oxylipins compared to general ALA amount in diatoms, PhytoPs are suspected to have a biological function (Lupette et al., 2018). Moreover, in plants, ALA serves as a primary precursor for various signaling compounds generated through oxidative modification by ROS (Ahme et al., 2020; Schaller and Stintzi, 2009). It would be thus interesting to lead further study on the possible implication of non-enzymatic oxylipins such as PhytoPs in migration response of MPB.

The deepoxydation rate also increased significantly between 250 and 500 PAR, indicating a high NPQ response (Doose and Hubas, 2024). In one hand, NPQ is considered to be the most crucial short-term photoacclimative processes in diatoms (Lavaud, 2007; Lavaud et al., 2002). These rapid responses can be activated within a few tens of seconds during a sudden rise in incident light intensity, so that they do not depend directly on gene regulation (Lavaud, 2007). On the other hand, the production of oxylipins such as jasmonate, is known to initiates an upregulation of antioxidant production through a feedback loop (Demmig-adams et al., 2012), notably by increasing concentrations of antioxidants such carotenoids. The xanthophylls are also known to directly scavenge the triplet-state excitation of Chl *a* (Larkum, 2003; Müller et al., 2001) and to have strong anti-oxidant properties. This, in turn, prevent lipid peroxidation and thus to inhibit additional oxylipin production (Andersson and Aro, 2001; Galinato et al., 2007; Havaux and Niyogi, 1999; Saniewski and Czapski, 1983; Wang and Zheng, 2005). It could be thus possible that the epoxidation or deepoxidation state of the xanthophylls influence isoprostanoids synthesis such the ALA and EPA derived oxylipins.

## 5 Conclusion

This study investigated for the first time the production of non-enzymatic oxylipins in a microphytobenthic biofilm under different light irradiances. It reveals that isoprostanoid levels in microphytobenthic biofilm respond to varying light intensities. They were identified to be originating from the diatoms, indicating a light-dependent influence on ROS production originating from the photosynthetic activity. The PUFAs precursors showed distinct peroxidation patterns under different light conditions, suggesting a link between light-induced ROS diversity, ROS scavenging efficiency by anti-oxidant system and PUFA oxidation pathways. Under 1000 PAR condition, the total amount of isoprotanoids increased indicating an oxidative stress. The EPA and AA derivatives characterized the HL conditions and evidenced lipid peroxidation, probably due to the anti-oxidants system becoming gradually overwhelmed face to the higher generation of ROS by the photosynthesis. On the contrary, the PhytoPs, ALA derivatives, characterized the LL and ML where the de-epoxidation state was low and ROS scavengers probably not overwhelmed. This indicated that the lipid peroxidation probably not occurred in the chloroplast but in others cellular sites such as endomembranes. The concomitant presence of the diatoms at the sediment surface and the PhytoP synthesis suggest that these oxygenic conditions could also partly influence this isoprostanoid production in a photosynthetic independent way. However, the PhytoPs resulted more likely from a regulated response of MPB organisms. These findings provide novel insights into oxylipin production in mudflat biofilms, highlighting the interest to explore their signaling roles related to photoprotective mechanisms and vertical migration.

## Author Contributions

Caroline Doose was in charge of designing and conducting the experiments, as well as the acquisition, analysis, and interpretation of data. Camille Oger contributed to the interpretation of the data and revised the work. Lindsay Mas-Normand conducted the extraction and measurement of the oxylipins, revised the work, and gave final approval for publication. Thierry Durand revised the work and gave final approval for publication. Cédric Hubas participated in the elaboration of the experimental design, data interpretation, and paper review.

## Reference

Ahme, O.S., Galano, J.M., Pavlickova, T., Revol-Cavalier, J., Vigor, C., Lee, J.C.Y., Oger, C., Durand, T., 2020. Moving forward with isoprostanes, neuroprostanes and phytoprostanes: Where are we now? Essays Biochem. 64, 463–484. 10.1042/EBC20190096

Améras, E., Stolz, S., Vollenweider, S., Reymond, P., Mène-Saffrané, L., Farmer, E.E., 2003. Reactive electrophile species activate defense gene expression in Arabidopsis. Plant J. 34, 205–216. 10.1046/J.1365-313X.2003.01718.X

Andersson, B., Aro, E.-M., 2001. Photodamage and D1 protein turnover in photosystem II. Regul. Photosynth. 377–393. 10.1007/0-306-48148-0_22

Apel, K., Hirt, H., 2004. Reactive oxygen species: Metabolism, oxidative stress, and signal transduction. Annu. Rev. Plant Biol. 55, 373–399. 10.1146/annurev.arplant.55.031903.141701

Asada, K., 1999. The water-water cycle in chloroplasts: Scavenging of active oxygens and dissipation of excess photons. Annu. Rev. Plant Biol. 50, 601–639.

Barnett, A., Méléder, V., Dupuy, C., Lavaud, J., 2020. The vertical migratory rhythm of intertidal microphytobenthos in sediment depends on the light photoperiod, intensity, and spectrum: evidence for a positive effect of blue wavelengths. Front. Mar. Sci. 7, 1–18. 10.3389/fmars.2020.00212

Beninger, P.G., Paterson, D.M., 2018. Mudflat Ecology, Aquatic Ec. ed. Springer Nature. 10.1007/978-3-319-99194-8

Büchel, C., Goss, R., Bailleul, B., Campbell, D.A., Lavaud, J., Lepetit, B., 2022. Photosynthetic light reactions in diatoms. I. The lipids and light-harvesting complexes of the thylakoid membrane. Mol. Life Diatoms 397–422. 10.1007/978-3-030-92499-7_15

Cartaxana, P., Ruivo, M., Hubas, C., Davidson, I., Serôdio, J., Jesus, B., 2011. Physiological versus behavioral photoprotection in intertidal epipelic and epipsammic benthic diatom communities. J. Exp. Mar. Bio. Ecol. 405, 120–127. 10.1016/J.JEMBE.2011.05.027

Consalvey, M., Paterson, D.M., Underwood, G.J.C., 2004. The ups and downs of life in a benthic biofilm: Migration of benthic diatoms. Diatom Res. 19, 181–202. 10.1080/0269249X.2004.9705870

Curien, G., Flori, S., Villanova, V., Magneschi, L., Giustini, C., Forti, G., Matringe, M., Petroutsos, D., Kuntz, M., Finazzi, G., 2016. The water to water cycles in microalgae. Plant Cell Physiol. 57, 1354–1363. 10.1093/PCP/PCW048

D’Autréaux, B., Toledano, M.B., 2007. ROS as signalling molecules: Mechanisms that generate specificity in ROS homeostasis. Nat. Rev. Mol. Cell Biol. 8, 813–824. 10.1038/nrm2256

D’Souza, F.M.L., Loneragan, N.R., 1999. Effects of monospecific and mixed-algae diets on survival, development and fatty acid composition of penaeid prawn (Penaeus spp.) larvae. Mar. Biol. 133, 621–633. 10.1007/S002270050502/METRICS

Dall’Osto, L., Cazzaniga, S., Havaux, M., Bassi, R., 2010. Enhanced photoprotection by protein-bound vs free xanthophyll pools: A comparative analysis of chlorophyll b and xanthophyll biosynthesis mutants. Mol. Plant 3, 576–593. 10.1093/mp/ssp117

Demmig-adams, B., Cohu, C.M., Zadelhoff, G., Veldink, G.A., Muller, O., Iii, W.W.A., 2012. Emerging trade-offs – impact of photoprotectants (PsbS, xanthophylls, and vitamin E) on oxylipins as regulators of development and defense. New Phytol.

Di Dato, V., Barbarinaldi, R., Amato, A., Di Costanzo, F., Fontanarosa, C., Perna, A., Amoresano, A., Esposito, F., Cutignano, A., Ianora, A., Romano, G., 2020a. Variation in prostaglandin metabolism during growth of the diatom Thalassiosira rotula. Sci. Reports 2020 101 10, 1–13. 10.1038/s41598-020-61967-3

Di Dato, V., Di Costanzo, F., Barbarinaldi, R., Perna, A., Ianora, A., Romano, G., 2019. Unveiling the presence of biosynthetic pathways for bioactive compounds in the Thalassiosira rotula transcriptome. Sci. Rep. 9, 1–15. 10.1038/s41598-019-46276-8

Di Dato, V., Ianora, A., Romano, G., 2020b. Identification of prostaglandin pathway in dinoflagellates by transcriptome data mining. Mar. Drugs 18, 1–17. 10.3390/MD18020109

Diaz, J.M., Hansel, C.M., Apprill, A., Brighi, C., Zhang, T., Weber, L., McNally, S., Xun, L., 2016. Species-specific control of external superoxide levels by the coral holobiont during a natural bleaching event. Nat. Commun. 2016 71 7, 1–10. 10.1038/ncomms13801

Diaz, J.M., Hansel, C.M., Voelker, B.M., Mendes, C.M., Andeer, P.F., Zhang, T., 2013. Widespread production of extracellular superoxide by heterotrophic bacteria. Science (80-.). 340, 1223–1226. 10.1126/science.1237331

Dietz, K.J., 2008. Redox signal integration: From stimulus to networks and genes. Physiol. Plant. 133, 459–468. 10.1111/j.1399-3054.2008.01120.x

Dixon, T.C., Vermilyea, A.W., Scott, D.T., Voelker, B.M., 2013. Hydrogen peroxide dynamics in an agricultural headwater stream: Evidence for significant nonphotochemical production. Limnol. Oceanogr. 58, 2133–2144. 10.4319/LO.2013.58.6.2133

Doose, C., Hubas, C., 2024. The metabolites of light: Untargeted metabolomic approaches bring new clues to understand light-driven acclimation of intertidal mudflat biofilm. Sci. Total Environ. 912. 10.1016/j.scitotenv.2023.168692

Dumanović, J., Nepovimova, E., Natić, M., Kuča, K., Jaćević, V., 2021. The significance of reactive oxygen species and antioxidant defense system in plants: A concise overview. Front. Plant Sci. 11, 1–13. 10.3389/FPLS.2020.552969/BIBTEX

Ezequiel, J., Laviale, M., Frankenbach, S., Cartaxana, P., Serôdio, J., 2015. Photoacclimation state determines the photobehaviour of motile microalgae: The case of a benthic diatom. J. Exp. Mar. Bio. Ecol. 468, 11–20. 10.1016/j.jembe.2015.03.004

Foyer, C.H., 2018. Reactive oxygen species, oxidative signaling and the regulation of photosynthesis. Environ. Exp. Bot. 154, 134–142. 10.1016/J.ENVEXPBOT.2018.05.003

Foyer, C.H., Ruban, A. V., Noctor, G., 2017. Viewing oxidative stress through the lens of oxidative signalling rather than damage. Biochem. J. 474, 877–883. 10.1042/BCJ20160814

Galano, J.M., Lee, Y.Y., Oger, C., Vigor, C., Vercauteren, J., Durand, T., Giera, M., Lee, J.C.Y., 2017. Isoprostanes, neuroprostanes and phytoprostanes: An overview of 25 years of research in chemistry and biology. Prog. Lipid Res. 68, 83–108. 10.1016/j.plipres.2017.09.004

Galinato, M.G.I., Niedzwiedzki, D., Deal, C., Birge, R.R., Frank, H.A., 2007. Cation radicals of xanthophylls. Photosynth. Res. 94, 67–78. 10.1007/s11120-007-9218-5

Gerwick, W.H., Moghaddam, M., Hamberg, M., 1991. Oxylipin metabolism in the red alga Gracilariopsis lemaneiformis: Mechanism of formation of vicinal dihydroxy fatty acids. Arch. Biochem. Biophys. 290, 436–444. 10.1016/0003-9861(91)90563-X

Guy, A., Oger, C., Heppekausen, J., Signorini, C., Defelice, C., Fürstner, A., Durand, T., Galano, J.M., 2014. Oxygenated Metabolites of n-3 polyunsaturated fatty acids as potential oxidative stress biomarkers: Total synthesis of 8-F3t-IsoP, 10-F 4t-NeuroP and [D4]-10-F4t-NeuroP. Chem. - A Eur. J. 20, 6374–6380. 10.1002/chem.201400380

Hansel, C.M., Diaz, J.M., Plummer, S., 2019. Tight regulation of extracellular superoxide points to its vital role in the physiology of the globally relevant Roseobacter clade. MBio 10. 10.1128/MBIO.02668-18

Haubois, A.G., Sylvestre, F., Guarini, J.M., Richard, P., Blanchard, G.F., 2005. Spatio-temporal structure of the epipelic diatom assemblage from an intertidal mudflat in Marennes-Oléron Bay, France. Estuar. Coast. Shelf Sci. 64, 385–394. 10.1016/j.ecss.2005.03.004

Havaux, M., Niyogi, K.K., 1999. The violaxanthin cycle protects plants from photooxidative damage by more than one mechanism. Proc. Natl. Acad. Sci. U. S. A. 96, 8762–8767. 10.1073/PNAS.96.15.8762/ASSET/925DD9F9-C3DC-4640-A8CB-DF689339A23E/ASSETS/GRAPHIC/PQ1690366004.JPEG

Hope, J.A., Paterson, D.M., Thrush, S.F., Julie Hope, C.A., Editor, H., Van Alstyne, K., 2020. The role of microphytobenthos in soft-sediment ecological networks and their contribution to the delivery of multiple ecosystem services. J. Ecol. 108, 815–830. 10.1111/1365-2745.13322

Ibrahim, A., Mbodji, K., Hassan, A., Aziz, M., Boukhettala, N., Coëffier, M., Savoye, G., Déchelotte, P., Marion-Letellier, R., 2011. Anti-inflammatory and anti-angiogenic effect of long chain n-3 polyunsaturated fatty acids in intestinal microvascular endothelium. Clin. Nutr. 30, 678–687. 10.1016/J.CLNU.2011.05.002

Jahn, U., Galano, J.M., Durand, T., 2008. Beyond prostaglandins - Chemistry and biology of cyclic oxygenated metabolites formed by free-radical pathways from polyunsaturated fatty acids. Angew. Chemie Int. Ed. 47, 5894–5955. 10.1002/anie.200705122

Jesus, B., Jauffrais, T., Trampe, E., Méléder, V., Ribeiro, L., Bernhard, J.M., Geslin, E., Kühl, M., 2023. Microscale imaging sheds light on species-specific strategies for photo-regulation and photo-acclimation of microphytobenthic diatoms. Environ. Microbiol. 1–17. 10.1111/1462-2920.16499

Jesus, B., Perkins, R.G., Consalvey, M., Brotas, V., Paterson, D.M., 2006. Effects of vertical migrations by benthic microalgae on fluorescence measurements of photophysiology. Mar. Ecol. Prog. Ser. 315, 55–66. 10.3354/meps315055

Kelly, J.R., Scheibling, R.E., 2012. Fatty acids as dietary tracers in benthic food webs. Mar. Ecol. Prog. Ser. 446, 1–22. 10.3354/meps09559

Knieper, M., Viehhauser, A., Dietz, K.-J., 2023. Oxylipins and reactive carbonyls as regulators of the plant redox and reactive oxygen species network under stress. Antioxidants 12, 814–829.

Krieger-Liszkay, A., Christian, A.E., Ae, F., Trebst, A., 2008. Singlet oxygen production in photosystem II and related protection mechanism. Photosynth. Res. 98, 551–564. 10.1007/s11120-008-9349-3

Lamari, N., Ruggiero, M.V., D’Ippolito, G., Kooistra, W.H.C.F., Fontana, A., Montresor, M., 2013. Specificity of lipoxygenase pathways supports species delineation in the marine diatom genus Pseudo-nitzschia. PLoS One 8, 1–10. 10.1371/JOURNAL.PONE.0073281

Larkum, A.W.D., 2003. Light-harvesting systems in algae, in: Larkum, W., Douglas, S., Raven, J. (Eds.), Photosynthesis in Algae. Kluwer Academic Publishers, pp. 277–304. 10.1007/978-94-007-1038-2_13

Lavaud, J., 2007. Fast regulation of photosynthesis in diatoms: mechanisms, evolution and ecophysiology.

Lavaud, J., Goss, R., 2014. The peculiar features of non-photochemical fluorescence quenching in diatoms and brown algae, in: Demmig-Adams, B., Garab, G., Adams III, W. Govindjee (Eds.), Non-photochemical quenching and energy dissipation in plants, algae and cyanobacteria. Springer Science+Business Media Dordrecht 2014, pp. 421–443. 10.1007/978-94-017-9032-1_20

Lavaud, J., Rousseau, B., Van Gorkom, H.J., Etienne, A.L., 2002. Influence of the diadinoxanthin pool size on photoprotection in the marine planktonic diatom Phaeodactylum tricornutum. Plant Physiol. 129, 1398–1406. 10.1104/PP.002014

Laviale, M., Frankenbach, S., Serôdio, J., 2016. The importance of being fast: comparative kinetics of vertical migration and non-photochemical quenching of benthic diatoms under light stress. Mar. Biol. 163, 1–12. 10.1007/s00227-015-2793-7

Learman, D.R., Voelker, B.M., Vazquez-Rodriguez, A.I., Hansel, C.M., 2011. Formation of manganese oxides by bacterially generated superoxide. Nat. Geosci. 4, 95–98. 10.1038/NGEO1055

Leblond, J.D., Dahmen, A.S., Dodson, V.J., Dahmen, J.L., 2013. Characterization of the betaine lipids, diacylglyceryl-N,N,N-trimethylhomoserine (DGTS) and diaclyglycerylhydroxymethyl-N,N,N-trimethyl-β-alanine (DGTA), in brown- and green-pigmented raphidophytes. Arch. Hydrobiol. Suppl. Algol. Stud. 142, 17–27. 10.1127/1864-1318/2013/0130

Lebreton, B., Rivaud, A., Picot, L., Prévost, B., Barillé, L., Sauzeau, T., Beseres Pollack, J., Lavaud, J., 2019. From ecological relevance of the ecosystem services concept to its socio-political use. The case study of intertidal bare mudflats in the Marennes-Oléron Bay, France. Ocean Coast. Manag. 172, 41–54. 10.1016/J.OCECOAMAN.2019.01.024

Lepetit, B., Sturm, S., Rogato, A., Gruber, A., Sachse, M., Falciatore, A., Kroth, P.G., Lavaud, J., 2013. High light acclimation in the secondary plastids containing diatom Phaeodactylum tricornutum is triggered by the redox state of the plastoquinone pool. Plant Physiol. 161, 853–865. 10.1104/PP.112.207811

Linares-Maurizi, A., Reversat, G., Awad, R., Bultel-Poncé, V., Oger, C., Galano, J.M., Balas, L., Durbec, A., Bertrand-Michel, J., Durand, T., Pradelles, R., Vigor, C., 2023. Bioactive oxylipins profile in marine microalgae. Mar. Drugs 21, 136. 10.3390/MD21030136/S1

Longini, M., Belvisi, E., Proietti, F., Bazzini, F., Buonocore, G., Perrone, S., 2017. Oxidative stress biomarkers: establishment of reference values for isoprostanes, AOPP, and NPBI in cord blood. Mediators Inflamm. 2017, 1–6. 10.1155/2017/1758432

Lupette, J., Jaussaud, A., Vigor, C., Oger, C., Galano, J.M., Réversat, G., Vercauteren, J., Jouhet, J., Durand, T., Maréchal, E., 2018. Non-enzymatic synthesis of bioactive isoprostanoids in the diatom Phaeodactylum following oxidative stress. Plant Physiol. 178, 1344–1357. 10.1104/PP.18.00925

Macintyre, H.L., Geider, R.J., Miller, D.C., 1996. Microphytobenthos: The ecological role of the “secret garden” of unvegetated, shallow-water marine habitats. I. Role in sediment stability and shallow-water food webs. Estuaries 19, 202–212. 10.2307/1352225

Mallick, N., Mohn, F.H., 2000. Reactive oxygen species: Response of algal cells. J. Plant Physiol. 157, 183–193. 10.1016/S0176-1617(00)80189-3

Marsico, R.M., Schneider, R.J., Voelker, B.M., Zhang, T., Diaz, J.M., Hansel, C.M., Ushijima, S., 2015. Spatial and temporal variability of widespread dark production and decay of hydrogen peroxide in freshwater. Aquat. Sci. 77, 523–533. 10.1007/S00027-015-0399-2/METRICS

Méléder, V., Rincé, Y., Barillé, L., Gaudin, P., Rosa, P., 2007. Spatiotemporal changes in microphytobenthos assemblages in a macrotidal flat (Bourgneuf Bay, France). J. Phycol. 43, 1177–1190. 10.1111/j.1529-8817.2007.00423.x

Meyer, A.J., 2008. The integration of glutathione homeostasis and redox signaling. J. Plant Physiol. 165, 1390–1403. 10.1016/j.jplph.2007.10.015

Meyer, N., Rettner, J., Werner, M., Werz, O., Pohnert, G., 2018. Algal oxylipins mediate the resistance of diatoms against algicidal bacteria. Mar. Drugs 16, 1–8. 10.3390/MD16120486

Mittler, R., 2017. ROS are good. Trends Plant Sci. 22, 11–19. 10.1016/j.tplants.2016.08.002

Mittler, R., Vanderauwera, S., Suzuki, N., Miller, G., Tognetti, V.B., Vandepoele, K., Gollery, M., Shulaev, V., Van Breusegem, F., 2011. ROS signaling: The new wave? Trends Plant Sci. 16, 300–309. 10.1016/j.tplants.2011.03.007

Montuschi, P., Barnes, P.J., Jackson Roberts Ii, L., 2004. Isoprostanes: markers and mediators of oxidative stress. FASEB J. 18, 1791–1800. 10.1096/FJ.04-2330REV

Morrow, J.D., Awad, J.A., Boss, H.J., Blair, I.A., Roberts, L.J., 1992. Non-cyclooxygenase-derived prostanoids (F2-isoprostanes) are formed in situ on phospholipids. Proc. Natl. Acad. Sci. U. S. A. 89, 10721–10725. 10.1073/PNAS.89.22.10721

Morrow, J.D., Hill, K.E., Burk, R.F., Nammour, T.M., Badr, K.F., Roberts, L.J., 1990. A series of prostaglandin F2-like compounds are produced in vivo in humans by a non-cyclooxygenase, free radical-catalyzed mechanism. Proc. Natl. Acad. Sci. U. S. A. 87, 9383–9387. 10.1073/PNAS.87.23.9383

Mueller, M.J., 2004. Archetype signals in plants: The phytoprostanes. Curr. Opin. Plant Biol. 7, 441–448. 10.1016/j.pbi.2004.04.001

Müller, P., Li, X.P., Niyogi, K.K., 2001. Non-photochemical quenching. A response to excess light energy. Plant Physiol. 125, 1558–1566. 10.1104/PP.125.4.1558

Mullineaux, P.M., Exposito-Rodriguez, M., Laissue, P.P., Smirnoff, N., 2018. ROS-dependent signalling pathways in plants and algae exposed to high light: Comparisons with other eukaryotes. Free Radic. Biol. Med. 122, 52–64. 10.1016/j.freeradbiomed.2018.01.033

Noctor, G., Foyer, C.H., 2016. Intracellular redox compartmentation and ROS-related communication in regulation and signaling. Plant Physiol. 171, 1581–1592. 10.1104/PP.16.00346

Nourooz-Zadeh, J., Liu, E.H.C., Änggård, E.E., Halliwell, B., 1998. F4-isoprostanes: A novel class of prostanoids formed during peroxidation of docosahexaenoic acid (DHA). Biochem. Biophys. Res. Commun. 242, 338–344. 10.1006/bbrc.1997.7883

Nymark, M., Valle, K.C., Brembu, T., Hancke, K., Winge, P., Andresen, K., Johnsen, G., Bones, A.M., 2009. An integrated analysis of molecular acclimation to high light in the marine diatom Phaeodactylum tricornutum. PLoS One 4, 1–14. 10.1371/journal.pone.0007743

Orefice, I., Di Dato, V., Sardo, A., Lauritano, C., Romano, G., 2022. Lipid mediators in marine diatoms. Aquat. Ecol. 377–397. 10.1007/s10452-021-09932-8

Parchmann, S., Mueller, M.J., 1998. Evidence for the formation of dinor isoprostanes E1 from alpha-linolenic acid in plants. J. Biol. Chem. 273, 32650–32655. 10.1074/JBC.273.49.32650

Parrish, C.C., 2013. Lipids in marine ecosystems. ISRN Oceanogr. 2013, 16.

Paul Hansard, S., Vermilyea, A.W., Voelker, B.M., 2010. Measurements of superoxide radical concentration and decay kinetics in the Gulf of Alaska. Deep. Res. Part I Oceanogr. Res. Pap. 57, 1111–1119. 10.1016/j.dsr.2010.05.007

Perkins, R.G., Lavaud, J., Serôdio, J., Mouget, J.L., Cartaxana, P., Rosa, P., Barille, L., Brotas, V., Jesus, B.M., 2010. Vertical cell movement is a primary response of intertidal benthic biofilms to increasing light dose. Mar. Ecol. Prog. Ser. 416, 93–103. 10.3354/meps08787

Pitzschke, A., Forzani, C., Hirt, H., 2006. Reactive oxygen species signaling in plants. Antioxid. Redox Signal. 8, 1757–1764. 10.1089/ARS.2006.8.1757

Prelle, L.R., Karsten, U., 2022. Photosynthesis, respiration, and growth of five benthic diatom strains as a function of intermixing processes of coastal peatlands with the baltic sea. Microorganisms 10, 1–20. 10.3390/microorganisms10040749

Prins, A., Deleris, P., Hubas, C., Jesus, B., 2020. Effect of light intensity and light quality on diatom behavioral and physiological photoprotection. Front. Mar. Sci. 7, 1–17. 10.3389/fmars.2020.00203

Ramel, F., Birtic, S., Ginies, C., Soubigou-Taconnat, L., Triantaphylidès, C., Havaux, M., 2012. Carotenoid oxidation products are stress signals that mediate gene responses to singlet oxygen in plants. Proc. Natl. Acad. Sci. U. S. A. 109, 5535–5540. 10.1073/PNAS.1115982109/SUPPL_FILE/SD01.XLS

Ribeiro, L., Brotas, V., Rincé, Y., Jesus, B., 2013. Structure and diversity of intertidal benthic diatom assemblages in contrasting shores: A case study from the Tagus estuary. J. Phycol. 49, 258–270. 10.1111/jpy.12031

Roberts, L.J., Montine, T.J., Markesbery, W.R., Tapper, A.R., Hardy, P., Chemtob, S., Dettbarn, W.D., Morrow, J.D., 1998. Formation of isoprostane-like compounds (neuroprostanes) in vivo from docosahexaenoic acid. J. Biol. Chem. 273, 13605–13612. 10.1074/JBC.273.22.13605

Roe, K.L., Schneider, R.J., Hansel, C.M., Voelker, B.M., 2016. Measurement of dark, particle-generated superoxide and hydrogen peroxide production and decay in the subtropical and temperate North Pacific Ocean. Deep Sea Res. Part I Oceanogr. Res. Pap. 107, 59–69. 10.1016/J.DSR.2015.10.012

Rose, A.L., Webb, E.A., Waite, T.D., Moffett, J.W., 2008. Measurement and implications of nonphotochemically generated superoxide in the equatorial pacific ocean. Environ. Sci. Technol. 42, 2387–2393. 10.1021/es7024609

Roy, J., Galano, J.M., Durand, T., Le Guennec, J.Y., Lee, J.C.Y., 2017. Physiological role of reactive oxygen species as promoters of natural defenses. FASEB J. 31, 3729–3745. 10.1096/FJ.201700170R

Ruocco, N., Albarano, L., Esposito, R., Zupo, V., Costantini, M., Ianora, A., 2020. Multiple roles of diatom-derived oxylipins within marine environments and their potential biotechnological applications. Mar. Drugs 18, 1–25. 10.3390/md18070342

Rusak, S.A., Peake, B.M., Richard, L.E., Nodder, S.D., Cooper, W.J., 2011. Distributions of hydrogen peroxide and superoxide in seawater east of New Zealand. Mar. Chem. 127, 155–169. 10.1016/j.marchem.2011.08.005

Russo, E., d’Ippolito, G., Fontana, A., Sarno, D., D’Alelio, D., Busseni, G., Ianora, A., von Elert, E., Carotenuto, Y., 2020. Density-dependent oxylipin production in natural diatom communities: possible implications for plankton dynamics. ISME J. 14, 164–177. 10.1038/S41396-019-0518-5

Saniewski, M., Czapski, J., 1983. The effect of methyl jasmonate on lycopene and β-carotene accumulation in ripening red tomatoes. Experientia 39, 1373–1374. 10.1007/BF01990110/METRICS

Schaller, A., Stintzi, A., 2009. Enzymes in jasmonate biosynthesis - structure, function, regulation. Phytochemistry 70, 1532–1538. 10.1016/J.PHYTOCHEM.2009.07.032

Schneider, R.J., Roe, K.L., Hansel, C.M., Voelker, B.M., 2016. Species-level variability in extracellular production rates of reactive oxygen species by diatoms. Front. Chem. 4, 5. 10.3389/fchem.2016.00005

Serôdio, J., Coelho, H., Vieira, S., Cruz, S., 2006. Microphytobenthos vertical migratory photoresponse as characterised by light-response curves of surface biomass. Estuar. Coast. Shelf Sci. 68, 547–556. 10.1016/j.ecss.2006.03.005

Serôdio, J., Ezequiel, J., Barnett, A., Mouget, J.L., Meĺéder, V., Laviale, M., Lavaud, J., 2012. Efficiency of photoprotection in microphytobenthos: role of vertical migration and the xanthophyll cycle against photoinhibition. Aquat. Microb. Ecol. 67, 161–175. 10.3354/AME01591

Serôdio, J., Vieira, S., Cruz, S., 2008. Photosynthetic activity, photoprotection and photoinhibition in intertidal microphytobenthos as studied in situ using variable chlorophyll fluorescence. Cont. Shelf Res. 28, 1363–1375. 10.1016/j.csr.2008.03.019

Sutherland, K.M., Coe, A., Gast, R.J., Plummer, S., Suffridge, C.P., Diaz, J.M., Bowman, J.S., Wankel, S.D., Hansel, C.M., 2019. Extracellular superoxide production by key microbes in the global ocean. Limnol. Oceanogr. 64, 2679–2693. 10.1002/lno.11247

Triantaphylidès, C., Havaux, M., 2009. Singlet oxygen in plants: production, detoxification and signaling. Trends Plant Sci. 14, 219–228. 10.1016/J.TPLANTS.2009.01.008

Triantaphylidès, C., Krischke, M., Hoeberichts, F.A., Ksas, B., Gresser, G., Havaux, M., Van Breusegem, F., Mueller, M.J., 2008. Singlet oxygen is the major reactive oxygen species involved in photooxidative damage to plants. Plant Physiol. 148, 960–968. 10.1104/PP.108.125690

Underwood, G.J.C., Kromkamp, J., 1999. Primary production by phytoplankton and microphytobenthos in estuaries. Adv. Ecol. Res. 29, 93–153. 10.1016/S0065-2504(08)60192-0

Vermilyea, A.W., Hansard, S.P., Voelker, B.M., 2010. Dark production of hydrogen peroxide in the Gulf of Alaska. Limnol. Oceanogr. 55, 580–588. 10.4319/LO.2010.55.2.0580

Vigor, C., Oger, C., Reversat, G., Rocher, A., Zhou, B., Linares-Maurizi, A., Guy, A., Bultel-Poncé, V., Galano, J.M., Vercauteren, J., Durand, T., Potin, P., Tonon, T., Leblanc, C., 2020. Isoprostanoid profiling of marine microalgae. Biomolecules 10, 1–20. 10.3390/BIOM10071073

Vigor, C., Reversat, G., Rocher, A., Oger, C., Vercauteren, J., Durand, T., Tonon, T., Potin, P., 2018. Isoprostanoids quantitative profiling of marine red and brown macroalgae. Food Chem. 268, 452–462. 10.1016/j.foodchem.2018.06.111

Vogelsang, L., Dietz, K.J., 2022. Plant thiol peroxidases as redox sensors and signal transducers in abiotic stress acclimation. Free Radic. Biol. Med. 193, 764–778. 10.1016/J.FREERADBIOMED.2022.11.019

Wang, S.Y., Zheng, W., 2005. Preharvest application of methyl jasmonate increases fruit quality and antioxidant capacity in raspberries. Int. J. Food Sci. Technol. 40, 187–195. 10.1111/J.1365-2621.2004.00930.X

Waring, J., Klenell, M., Bechtold, U., Underwood, G.J.C., Baker, N.R., 2010. Light-induced responses of oxygen photoreduction, reactive oxygen species production and scavenging in two diatom species. J. Phycol. 46, 1206–1217. 10.1111/j.1529-8817.2010.00919.x

Werner, U., Billerbeck, M., Polerecky, L., Franke, U., Huettel, M., Van Beusekom, J.E.E., De Beer, D., 2006. Spatial and temporal patterns of mineralization rates and oxygen distribution in a permeable intertidal sand flat (Sylt, Germany). Limnol. Oceanogr. 51, 2549–2563. 10.4319/lo.2006.51.6.2549

Woelfel, J., Schoknecht, A., Schaub, I., Enke, N., Schumann, R., Karsten, U., 2014. Growth and photosynthesis characteristics of three benthic diatoms from the brackish southern Baltic Sea in relation to varying environmental conditions. Phycologia 53, 639–651. 10.2216/14-019.1

Zhang, T., Diaz, J.M., Brighi, C., Parsons, R.J., McNally, S., Apprill, A., Hansel, C.M., 2016. Dark production of extracellular superoxide by the coral Porites astreoides and representative symbionts. Front. Mar. Sci. 3, 1–16. 10.3389/FMARS.2016.00232/BIBTEX

