## Supplementary material for "Non-enzymatic oxylipin production in a mudflat microphytobenthic biofilm: evidence of a diatom response to light": Table S1

**Table S1.** *Concentration of the oxylipins in pg/mg dw of biofilm measured in biofilm exposed to dark and to irradiances of low lights (LL: 50, 100), medium light (ML: 250) and high lights (HL: 500, 750, 1000) in µmol photons $m^{-2}$ $s^{-1}$PAR.*

| Light | Oxylipines | Family | Precursor | Concentration | SD |
|---|---|---|---|---|---|
| D | 10-*epi*-10-$F_{4t}$-NeuroP | NeuroP | DHA | 4,0 | 1,2 |
| D | 10-$F_{4t}$-NeuroP | NeuroP | DHA | 2,1 | 0,8 |
| D | 14-*epi*-14-$F_{4t}$-NeuroP | NeuroP | DHA | 1,8 | 0,7 |
| D | 14-$F_{4t}$-NeuroP | NeuroP | DHA | 2,6 | 0,8 |
| D | 15(*R*)-$PGF_2$ | Prostaglandin | AA | 3,0 | 0,5 |
| D | 15-*epi*-15-$F_{2t}$-IsoP | IsoP | AA | 3,4 | 0,6 |
| D | 15-$F_{2t}$-IsoP | IsoP | AA | 1,7 | 0,2 |
| D | 16-$B_{1t}$-PhytoP | PhytoP | ALA | 1,5 | 0,5 |
| D | 18-*epi*-18-$F_{3t}$-IsoP | IsoP | EPA | 29,7 | 6,0 |
| D | 18-$F_{3t}$-IsoP | IsoP | EPA | 11,5 | 1,9 |
| D | 20-*epi*--20-$F_{4t}$-NeuroP | NeuroP | DHA | 4,1 | 1,0 |
| D | 20-$F_{4t}$-NeuroP | NeuroP | DHA | 4,0 | 1,2 |
| D | 4(*RS*)-4-$F_{4t}$-NeuroP | NeuroP | DHA | 4,4 | 0,8 |
| D | 5-$F_{2c}$-IsoP | IsoP | AA | 10,9 | 0,8 |
| D | 5-*epi*-5-$F_{3t}$-IsoP | IsoP | EPA | 121,7 | 15,7 |
| D | 5(*RS*)-5-$F_{2t}$-IsoP | IsoP | AA | 4,0 | 0,6 |
| D | 5-$F_{3t}$-IsoP | IsoP | EPA | 37,8 | 5,0 |
| D | 8-*epi*-8-$F_{3t}$-IsoP | IsoP | EPA | 4,0 | 0,6 |
| D | 8-$F_{3t}$-IsoP | IsoP | EPA | 3,0 | 0,6 |
| D | 9-*epi*-9-$F_{1t}$-PhytoP + *ent*-16-$F_{1t}$-PhytoP | PhytoP | ALA | 38,7 | 8,9 |
| D | 9-$F_{1t}$-PhytoP | PhytoP | ALA | 29,1 | 7,1 |
| D | 9-$L_{1t}$-PhytoP | PhytoP | ALA | 1,4 | 0,5 |
| D | *ent*-16-A-9-*epi*-ST-$\Delta^{14}$-10-PhytoF | PhytoF | ALA | 1,0 | 0,4 |
| D | *ent*-16-B-9-*epi*-ST-$\Delta^{14}$-10-PhytoF | PhytoF | ALA | 1,5 | 0,9 |
| D | *ent*-16-*epi*-16-$F_{1t}$-PhytoP | PhytoP | ALA | 14,0 | 3,6 |
| D | *ent*-16-$F_{1t}$-PhytoP + 9-*epi*-9-$F_{1t}$-PhytoP | PhytoP | ALA | 38,7 | 8,9 |
| D | *ent*-7(*RS*)-7-$F_{2t}$-dihomo-IsoP | IsoP | AdA | 1,1 | 0,2 |
| D | $PGF_2$ | Prostaglandin | AA | 3,7 | 0,8 |
| D | $PGF_3$ | Prostaglandin | EPA | 36,8 | 7,6 |
| 50 | 10-*epi*-10-$F_{4t}$-NeuroP | NeuroP | DHA | 4,2 | 1,1 |
| 50 | 10-$F_{4t}$-NeuroP | NeuroP | DHA | 2,6 | 1,1 |
| 50 | 14-*epi*-14-$F_{4t}$-NeuroP | NeuroP | DHA | 2,2 | 0,8 |
| 50 | 14-$F_{4t}$-NeuroP | NeuroP | DHA | 3,0 | 0,9 |
| 50 | 15(*R*)-$PGF_2$ | Prostaglandin | AA | 3,1 | 0,4 |
| 50 | 15-*epi*-15-$F_{2t}$-IsoP | IsoP | AA | 3,3 | 0,4 |
| 50 | 15-$F_{2t}$-IsoP | IsoP | AA | 1,8 | 0,3 |
| 50 | 16-$B_{1t}$-PhytoP | PhytoP | ALA | 1,7 | 0,7 |
| 50 | 18-*epi*-18-$F_{3t}$-IsoP | IsoP | EPA | 27,1 | 4,0 |
| 50 | 18-$F_{3t}$-IsoP | IsoP | EPA | 12,8 | 2,1 |
| 50 | 20-*epi*--20-$F_{4t}$-NeuroP | NeuroP | DHA | 4,0 | 1,0 |
| 50 | 20-$F_{4t}$-NeuroP | NeuroP | DHA | 4,3 | 1,5 |
| 50 | 4(*RS*)-4-$F_{4t}$-NeuroP | NeuroP | DHA | 4,9 | 1,0 |
| 50 | 5-$F_{2c}$-IsoP | IsoP | AA | 11,2 | 1,8 |
| 50 | 5-*epi*-5-$F_{3t}$-IsoP | IsoP | EPA | 126,4 | 17,3 |
| 50 | 5(*RS*)-5-$F_{2t}$-IsoP | IsoP | AA | 4,1 | 0,5 |
| 50 | 5-$F_{3t}$-IsoP | IsoP | EPA | 41,4 | 6,9 |
| 50 | 8-*epi*-8-$F_{3t}$-IsoP | IsoP | EPA | 4,9 | 0,9 |
| 50 | 8-$F_{3t}$-IsoP | IsoP | EPA | 3,2 | 0,7 |
| 50 | 9-*epi*-9-$F_{1t}$-PhytoP + *ent*-16-$F_{1t}$-PhytoP | PhytoP | ALA | 43,0 | 4,0 |
| 50 | 9-$F_{1t}$-PhytoP | PhytoP | ALA | 32,4 | 3,8 |
| 50 | 9-$L_{1t}$-PhytoP | PhytoP | ALA | 1,6 | 0,7 |
| 50 | *ent*-16-A-9-*epi*-ST-$\Delta^{14}$-10-PhytoF | PhytoF | ALA | 1,1 | 0,5 |
| 50 | *ent*-16-B-9-*epi*-ST-$\Delta^{14}$-10-PhytoF | PhytoF | ALA | 1,7 | 0,8 |
| 50 | *ent*-16-*epi*-16-$F_{1t}$-PhytoP | PhytoP | ALA | 15,7 | 1,4 |
| 50 | *ent*-16-$F_{1t}$-PhytoP + 9-*epi*-9-$F_{1t}$-PhytoP | PhytoP | ALA | 43,0 | 4,0 |
| 50 | *ent*-7(*RS*)-7-$F_{2t}$-dihomo-IsoP | IsoP | AdA | 1,3 | 0,2 |
| 50 | $PGF_2$ | Prostaglandin | AA | 4,5 | 0,5 |
| 50 | $PGF_3$ | Prostaglandin | EPA | 40,1 | 8,5 |
| 100 | 10-*epi*-10-$F_{4t}$-NeuroP | NeuroP | DHA | 3,5 | 0,6 |
| 100 | 10-$F_{4t}$-NeuroP | NeuroP | DHA | 1,9 | 0,3 |
| 100 | 14-*epi*-14-$F_{4t}$-NeuroP | NeuroP | DHA | 1,8 | 0,7 |
| 100 | 14-$F_{4t}$-NeuroP | NeuroP | DHA | 2,1 | 0,3 |
| 100 | 15(*R*)-$PGF_2$ | Prostaglandin | AA | 3,4 | 0,8 |
| 100 | 15-*epi*-15-$F_{2t}$-IsoP | IsoP | AA | 3,4 | 0,9 |
| 100 | 15-$F_{2t}$-IsoP | IsoP | AA | 1,9 | 0,4 |
| 100 | 16-$B_{1t}$-PhytoP | PhytoP | ALA | 1,3 | 0,4 |
| 100 | 18-*epi*-18-$F_{3t}$-IsoP | IsoP | EPA | 28,4 | 5,2 |
| 100 | 18-$F_{3t}$-IsoP | IsoP | EPA | 13,1 | 2,2 |
| 100 | 20-*epi*--20-$F_{4t}$-NeuroP | NeuroP | DHA | 4,2 | 0,9 |
| 100 | 20-$F_{4t}$-NeuroP | NeuroP | DHA | 3,7 | 0,8 |

| | | | | | |
|---|---|---|---|---|---|
| 100 | 4(*RS*)-4-$F_{4t}$-NeuroP | NeuroP | DHA | 4,6 | 0,5 |
| 100 | 5-$F_{2c}$-IsoP | IsoP | AA | 13,3 | 2,5 |
| 100 | 5-*epi*-5-$F_{3t}$-IsoP | IsoP | EPA | 141,6 | 21,2 |
| 100 | 5(*RS*)-5-$F_{2t}$-IsoP | IsoP | AA | 4,5 | 0,7 |
| 100 | 5-$F_{3t}$-IsoP | IsoP | EPA | 44,8 | 5,4 |
| 100 | 8-*epi*-8-$F_{3t}$-IsoP | IsoP | EPA | 5,1 | 0,7 |
| 100 | 8-$F_{3t}$-IsoP | IsoP | EPA | 3,4 | 0,5 |
| 100 | 9-*epi*-9-$F_{1t}$-PhytoP + *ent*-16-$F_{1t}$-PhytoP | PhytoP | ALA | 46,9 | 10,1 |
| 100 | 9-$F_{1t}$-PhytoP | PhytoP | ALA | 35,3 | 7,7 |
| 100 | 9-$L_{1t}$-PhytoP | PhytoP | ALA | 1,1 | 0,4 |
| 100 | *ent*-16-A-9-*epi*-ST-$\Delta^{14}$-10-PhytoF | PhytoF | ALA | 0,9 | 0,3 |
| 100 | *ent*-16-B-9-*epi*-ST-$\Delta^{14}$-10-PhytoF | PhytoF | ALA | 1,4 | 0,7 |
| 100 | *ent*-16-*epi*-16-$F_{1t}$-PhytoP | PhytoP | ALA | 17,2 | 3,6 |
| 100 | *ent*-16-$F_{1t}$-PhytoP + 9-*epi*-9-$F_{1t}$-PhytoP | PhytoP | ALA | 46,9 | 10,1 |
| 100 | *ent*-7(*RS*)-7-$F_{2t}$-dihomo-IsoP | IsoP | AdA | 1,6 | 0,3 |
| 100 | $PGF_2$ | Prostaglandin | AA | 5,6 | 1,6 |
| 100 | $PGF_3$ | Prostaglandin | EPA | 48,0 | 10,0 |
| 250 | 10-*epi*-10-$F_{4t}$-NeuroP | NeuroP | DHA | 4,6 | 3,8 |
| 250 | 10-$F_{4t}$-NeuroP | NeuroP | DHA | 2,9 | 3,1 |
| 250 | 14-*epi*-14-$F_{4t}$-NeuroP | NeuroP | DHA | 2,3 | 2,5 |
| 250 | 14-$F_{4t}$-NeuroP | NeuroP | DHA | 3,1 | 2,9 |
| 250 | 15(*R*)-$PGF_2$ | Prostaglandin | AA | 3,5 | 0,2 |
| 250 | 15-*epi*-15-$F_{2t}$-IsoP | IsoP | AA | 2,9 | 0,3 |
| 250 | 15-$F_{2t}$-IsoP | IsoP | AA | 1,6 | 0,2 |
| 250 | 16-$B_{1t}$-PhytoP | PhytoP | ALA | 1,2 | 0,6 |
| 250 | 18-*epi*-18-$F_{3t}$-IsoP | IsoP | EPA | 26,7 | 5,5 |
| 250 | 18-$F_{3t}$-IsoP | IsoP | EPA | 12,6 | 2,0 |
| 250 | 20-*epi*--20-$F_{4t}$-NeuroP | NeuroP | DHA | 4,8 | 3,5 |
| 250 | 20-$F_{4t}$-NeuroP | NeuroP | DHA | 3,7 | 2,3 |
| 250 | 4(*RS*)-4-$F_{4t}$-NeuroP | NeuroP | DHA | 5,1 | 3,3 |
| 250 | 5-$F_{2c}$-IsoP | IsoP | AA | 11,1 | 1,0 |
| 250 | 5-*epi*-5-$F_{3t}$-IsoP | IsoP | EPA | 129,6 | 14,2 |
| 250 | 5(*RS*)-5-$F_{2t}$-IsoP | IsoP | AA | 3,7 | 0,2 |
| 250 | 5-$F_{3t}$-IsoP | IsoP | EPA | 39,1 | 6,5 |
| 250 | 8-*epi*-8-$F_{3t}$-IsoP | IsoP | EPA | 5,0 | 1,7 |
| 250 | 8-$F_{3t}$-IsoP | IsoP | EPA | 3,5 | 1,7 |
| 250 | 9-*epi*-9-$F_{1t}$-PhytoP + *ent*-16-$F_{1t}$-PhytoP | PhytoP | ALA | 44,5 | 6,3 |
| 250 | 9-$F_{1t}$-PhytoP | PhytoP | ALA | 34,2 | 4,6 |
| 250 | 9-$L_{1t}$-PhytoP | PhytoP | ALA | 1,1 | 0,6 |
| 250 | *ent*-16-A-9-*epi*-ST-$\Delta^{14}$-10-PhytoF | PhytoF | ALA | 0,7 | 0,1 |
| 250 | *ent*-16-B-9-*epi*-ST-$\Delta^{14}$-10-PhytoF | PhytoF | ALA | 1,1 | 0,2 |
| 250 | *ent*-16-*epi*-16-$F_{1t}$-PhytoP | PhytoP | ALA | 16,4 | 2,2 |
| 250 | *ent*-16-$F_{1t}$-PhytoP + 9-*epi*-9-$F_{1t}$-PhytoP | PhytoP | ALA | 44,5 | 6,3 |
| 250 | *ent*-7(*RS*)-7-$F_{2t}$-dihomo-IsoP | IsoP | AdA | 1,2 | 0,2 |
| 250 | $PGF_2$ | Prostaglandin | AA | 4,9 | 0,8 |
| 250 | $PGF_3$ | Prostaglandin | EPA | 44,4 | 13,9 |
| 500 | 10-*epi*-10-$F_{4t}$-NeuroP | NeuroP | DHA | 3,3 | 0,6 |
| 500 | 10-$F_{4t}$-NeuroP | NeuroP | DHA | 1,5 | 0,2 |
| 500 | 14-*epi*-14-$F_{4t}$-NeuroP | NeuroP | DHA | 1,2 | 0,3 |
| 500 | 14-$F_{4t}$-NeuroP | NeuroP | DHA | 1,6 | 0,2 |
| 500 | 15(*R*)-$PGF_2$ | Prostaglandin | AA | 3,7 | 0,2 |
| 500 | 15-*epi*-15-$F_{2t}$-IsoP | IsoP | AA | 3,1 | 0,1 |
| 500 | 15-$F_{2t}$-IsoP | IsoP | AA | 1,7 | 0,1 |
| 500 | 16-$B_{1t}$-PhytoP | PhytoP | ALA | 1,4 | 0,4 |
| 500 | 18-*epi*-18-$F_{3t}$-IsoP | IsoP | EPA | 30,6 | 2,8 |
| 500 | 18-$F_{3t}$-IsoP | IsoP | EPA | 13,1 | 1,6 |
| 500 | 20-*epi*--20-$F_{4t}$-NeuroP | NeuroP | DHA | 3,7 | 0,8 |
| 500 | 20-$F_{4t}$-NeuroP | NeuroP | DHA | 3,3 | 0,9 |
| 500 | 4(*RS*)-4-$F_{4t}$-NeuroP | NeuroP | DHA | 3,7 | 0,7 |
| 500 | 5-$F_{2c}$-IsoP | IsoP | AA | 11,8 | 1,1 |
| 500 | 5-*epi*-5-$F_{3t}$-IsoP | IsoP | EPA | 140,8 | 23,7 |
| 500 | 5(*RS*)-5-$F_{2t}$-IsoP | IsoP | AA | 3,8 | 0,6 |
| 500 | 5-$F_{3t}$-IsoP | IsoP | EPA | 38,3 | 6,0 |
| 500 | 8-*epi*-8-$F_{3t}$-IsoP | IsoP | EPA | 5,2 | 0,2 |
| 500 | 8-$F_{3t}$-IsoP | IsoP | EPA | 3,7 | 0,4 |
| 500 | 9-*epi*-9-$F_{1t}$-PhytoP + *ent*-16-$F_{1t}$-PhytoP | PhytoP | ALA | 31,3 | 4,6 |
| 500 | 9-$F_{1t}$-PhytoP | PhytoP | ALA | 24,3 | 3,8 |
| 500 | 9-$L_{1t}$-PhytoP | PhytoP | ALA | 1,2 | 0,4 |
| 500 | *ent*-16-A-9-*epi*-ST-$\Delta^{14}$-10-PhytoF | PhytoF | ALA | 0,9 | 0,1 |
| 500 | *ent*-16-B-9-*epi*-ST-$\Delta^{14}$-10-PhytoF | PhytoF | ALA | 1,5 | 0,3 |
| 500 | *ent*-16-*epi*-16-$F_{1t}$-PhytoP | PhytoP | ALA | 11,9 | 2,0 |
| 500 | *ent*-16-$F_{1t}$-PhytoP + 9-*epi*-9-$F_{1t}$-PhytoP | PhytoP | ALA | 31,3 | 4,6 |
| 500 | *ent*-7(*RS*)-7-$F_{2t}$-dihomo-IsoP | IsoP | AdA | 0,9 | 0,1 |
| 500 | $PGF_2$ | Prostaglandin | AA | 5,4 | 1,1 |
| 500 | $PGF_3$ | Prostaglandin | EPA | 46,7 | 6,6 |
| 750 | 10-*epi*-10-$F_{4t}$-NeuroP | NeuroP | DHA | 3,6 | 0,5 |
| 750 | 10-$F_{4t}$-NeuroP | NeuroP | DHA | 1,7 | 0,1 |
| 750 | 14-*epi*-14-$F_{4t}$-NeuroP | NeuroP | DHA | 1,3 | 0,2 |
| 750 | 14-$F_{4t}$-NeuroP | NeuroP | DHA | 2,0 | 0,6 |

| | | | | | |
|---|---|---|---|---|---|
| 750 | 15(*R*)-$PGF_2$ | Prostaglandin | AA | 3,7 | 0,5 |
| 750 | 15-*epi*-15-$F_{2t}$-IsoP | IsoP | AA | 3,0 | 0,4 |
| 750 | 15-$F_{2t}$-IsoP | IsoP | AA | 1,7 | 0,2 |
| 750 | 16-$B_{1t}$-PhytoP | PhytoP | ALA | 1,1 | 0,1 |
| 750 | 18-*epi*-18-$F_{3t}$-IsoP | IsoP | EPA | 29,8 | 4,2 |
| 750 | 18-$F_{3t}$-IsoP | IsoP | EPA | 14,3 | 2,1 |
| 750 | 20-*epi*--20-$F_{4t}$-NeuroP | NeuroP | DHA | 4,1 | 0,4 |
| 750 | 20-$F_{4t}$-NeuroP | NeuroP | DHA | 3,5 | 0,6 |
| 750 | 4(*RS*)-4-$F_{4t}$-NeuroP | NeuroP | DHA | 4,3 | 0,6 |
| 750 | 5-$F_{2c}$-IsoP | IsoP | AA | 11,3 | 1,4 |
| 750 | 5-*epi*-5-$F_{3t}$-IsoP | IsoP | EPA | 153,9 | 26,0 |
| 750 | 5(*RS*)-5-$F_{2t}$-IsoP | IsoP | AA | 3,6 | 0,6 |
| 750 | 5-$F_{3t}$-IsoP | IsoP | EPA | 43,2 | 6,0 |
| 750 | 8-*epi*-8-$F_{3t}$-IsoP | IsoP | EPA | 4,9 | 0,7 |
| 750 | 8-$F_{3t}$-IsoP | IsoP | EPA | 3,8 | 0,6 |
| 750 | 9-*epi*-9-$F_{1t}$-PhytoP + *ent*-16-$F_{1t}$-PhytoP | PhytoP | ALA | 30,1 | 4,6 |
| 750 | 9-$F_{1t}$-PhytoP | PhytoP | ALA | 23,8 | 4,2 |
| 750 | 9-$L_{1t}$-PhytoP | PhytoP | ALA | 0,9 | 0,1 |
| 750 | *ent*-16-A-9-*epi*-ST-$\Delta^{14}$-10-PhytoF | PhytoF | ALA | 0,8 | 0,1 |
| 750 | *ent*-16-B-9-*epi*-ST-$\Delta^{14}$-10-PhytoF | PhytoF | ALA | 1,2 | 0,2 |
| 750 | *ent*-16-*epi*-16-$F_{1t}$-PhytoP | PhytoP | ALA | 11,6 | 1,9 |
| 750 | *ent*-16-$F_{1t}$-PhytoP + 9-*epi*-9-$F_{1t}$-PhytoP | PhytoP | ALA | 30,1 | 4,6 |
| 750 | *ent*-7(*RS*)-7-$F_{2t}$-dihomo-IsoP | IsoP | AdA | 1,1 | 0,1 |
| 750 | $PGF_2$ | Prostaglandin | AA | 5,4 | 1,0 |
| 750 | $PGF_3$ | Prostaglandin | EPA | 51,0 | 5,9 |
| 1000 | 10-*epi*-10-$F_{4t}$-NeuroP | NeuroP | DHA | 3,8 | 0,2 |
| 1000 | 10-$F_{4t}$-NeuroP | NeuroP | DHA | 2,0 | 0,1 |
| 1000 | 14-*epi*-14-$F_{4t}$-NeuroP | NeuroP | DHA | 1,7 | 0,2 |
| 1000 | 14-$F_{4t}$-NeuroP | NeuroP | DHA | 2,5 | 0,4 |
| 1000 | 15(*R*)-$PGF_2$ | Prostaglandin | AA | 3,8 | 0,5 |
| 1000 | 15-*epi*-15-$F_{2t}$-IsoP | IsoP | AA | 3,5 | 0,1 |
| 1000 | 15-$F_{2t}$-IsoP | IsoP | AA | 1,7 | 0,2 |
| 1000 | 16-$B_{1t}$-PhytoP | PhytoP | ALA | 1,4 | 0,2 |
| 1000 | 18-*epi*-18-$F_{3t}$-IsoP | IsoP | EPA | 38,9 | 1,8 |
| 1000 | 18-$F_{3t}$-IsoP | IsoP | EPA | 16,8 | 0,7 |
| 1000 | 20-*epi*--20-$F_{4t}$-NeuroP | NeuroP | DHA | 4,5 | 0,9 |
| 1000 | 20-$F_{4t}$-NeuroP | NeuroP | DHA | 4,7 | 0,6 |
| 1000 | 4(*RS*)-4-$F_{4t}$-NeuroP | NeuroP | DHA | 5,0 | 0,6 |
| 1000 | 5-$F_{2c}$-IsoP | IsoP | AA | 13,7 | 1,7 |
| 1000 | 5-*epi*-5-$F_{3t}$-IsoP | IsoP | EPA | 184,3 | 13,2 |
| 1000 | 5(*RS*)-5-$F_{2t}$-IsoP | IsoP | AA | 4,5 | 0,6 |
| 1000 | 5-$F_{3t}$-IsoP | IsoP | EPA | 51,8 | 4,4 |
| 1000 | 8-*epi*-8-$F_{3t}$-IsoP | IsoP | EPA | 5,6 | 0,5 |
| 1000 | 8-$F_{3t}$-IsoP | IsoP | EPA | 4,5 | 0,2 |
| 1000 | 9-*epi*-9-$F_{1t}$-PhytoP + *ent*-16-$F_{1t}$-PhytoP | PhytoP | ALA | 25,3 | 3,1 |
| 1000 | 9-$F_{1t}$-PhytoP | PhytoP | ALA | 19,7 | 1,9 |
| 1000 | 9-$L_{1t}$-PhytoP | PhytoP | ALA | 1,2 | 0,2 |
| 1000 | *ent*-16-A-9-*epi*-ST-$\Delta^{14}$-10-PhytoF | PhytoF | ALA | 0,9 | 0,2 |
| 1000 | *ent*-16-B-9-*epi*-ST-$\Delta^{14}$-10-PhytoF | PhytoF | ALA | 1,3 | 0,3 |
| 1000 | *ent*-16-*epi*-16-$F_{1t}$-PhytoP | PhytoP | ALA | 9,8 | 1,0 |
| 1000 | *ent*-16-$F_{1t}$-PhytoP + 9-*epi*-9-$F_{1t}$-PhytoP | PhytoP | ALA | 25,3 | 3,1 |
| 1000 | *ent*-7(*RS*)-7-$F_{2t}$-dihomo-IsoP | IsoP | AdA | 1,3 | 0,3 |
| 1000 | $PGF_2$ | Prostaglandin | AA | 5,4 | 0,6 |
| 1000 | $PGF_3$ | Prostaglandin | EPA | 64,1 | 3,7 |
